# DNA Sequence Differentially Modulates Histone Core and Tail Interactions in Canonical and Centromeric Nucleosomes

**DOI:** 10.64898/2026.11.12.766163

**Authors:** Abhik Ghosh Moulick, Sylvia Erhardt, Mariana Kozlowska

## Abstract

Human centromeric chromatin is a specialized chromatin domain characterized by CENP-A-containing nucleosomes interspersed with canonical histone H3 nucleosomes. It is assembled on tandem repeats of *α*-satellite DNA and defines centromere identity. How histone variant identity and DNA sequence jointly shape nucleosome structure and histone-DNA interactions that underpin centromeric chromatin formation remains incompletely understood. Here, we investigate the structural plasticity of the histone core and nucleosomal DNA, histone tail dynamics and DNA-sequence dependent changes for canonical (H3) and centromeric (CENP-A) nucleosomes using the SIRAH coarse-grained molecular dynamics (CG-MD) simulations. Nucleosomes containing *α*-satellite positioning sequence (ASP) and Widom 601 DNA sequence are systematically analyzed and compared. Our simulations reveal that the centromeric nucleosome exhibits a more stable histone core with lower structural plasticity and the histone tails engage DNA differently with an overall reduction in the histone-DNA contacts compared to the canonical nucleosome. Further investigation of CENP-N binding to the CENP-A nucleosome demonstrates substantial modulation of nucleosomal DNA dynamics and histone-DNA interactions. Replacing the ASP DNA sequence with the Widom 601 sequence preserves the stability of both types of the histone core while markedly altering histone-tail dynamics. The conformational dynamics of canonical nucleosomes is more strongly affected by the Widom 601 DNA sequence than those of their centromeric counterparts. However, CENP-N binding induces more spatially extensive changes in DNA geometry in Widom 601 than in ASP DNA. Our findings suggest that DNA sequence and histone composition have residue-level differences translating into distinct nucleosome dynamics, which could be among the determinants governing the structural and dynamic properties of centromeric chromatin and its biological function.

**SIGNIFICANCE:** Centromeric chromatin is essential for accurate chromosome segregation. How DNA sequence shapes the dynamics and histone-DNA interactions of nucleosomes, with consequences for centromeric chromatin folding, remains unclear. Using molecular dynamics simulations, we reveal that nucleosomes with CENP-A histone exhibit reduced histone core structural plasticity, and distinct sequence-dependent differences in DNA geometry and histone tail-DNA interactions in comparison to canonical H3 nucleosomes. Compared with α-satellite DNA, Widom 601 DNA increases the distance between the DNA gyres in CENP-A nucleosomes while having little effect in H3 nucleosomes. Binding of a centromere associated protein CENP-N induces sequence-dependent local DNA rear-rangements and causes a change of CENP-A-DNA contacts with Widom 601. These findings show that the choice of DNA sequence can attenuate histone-dependent differences in nucleosome geometry and alter the local response to CENP-N, highlighting its importance when interpreting centromeric nucleosome dynamics.

## INTRODUCTION

CENP-A is a histone H3 variant specifically enriched in nucleosomes within centromeric chromatin. In humans, this chromatin is typically assembled on repetitive *α*-satellite DNA sequences, where CENP-A coexists with canonical nucleosomes containing H3 histone. Quantitatively, CENP-A-containing nucleosomes represent only a small fraction of nucleosomes within the predominantly H3-based nucleosomes at centromeric chromatin (1). These otherwise sparsely distributed CENP-A nucleosomes are concentrated within hypomethylated centromere dip regions (CDRs), where the local ratio reaches approximately one CENP-A nucleosome for every four canonical nucleosomes (2, 3). Despite their relatively low overall abundance, CENP-A nucleosomes define centromere identity and provide a platform for kinetochore assembly, supporting accurate chromosome segregation (4, 5). However, the higher order folding and three dimensional architecture of CENP-A containing chromatin remain poorly understood, limiting our molecular-level understanding of its biological function.

The functional distinction between centromeric and canonical nucleosomes has a defined structural origin. CENP-A and H3 differ most substantially in the CENP-A targeting domain (CATD) of CENP-A, which is necessary and sufficient to confer centromeric identity (6). Crystal structures of human CENP-A nucleosome revealed a shortened *α*N helix, leaving approximately 13 bp of DNA at each terminus loosely held relative to the canonical particle (7). These features are the structural basis for the altered dynamics reported in subsequent biophysical and computational works(8–12), and they make the CENP-A nucleosome a plausible point at which DNA sequence could exert a differential effect. Furthermore, considering that centromeric chromatin contains both CENP-A- and H3-containing nucleosomes, understanding their structural and dynamical differences is important for identifying the molecular determinants that may contribute to chromatin folding, maintenance, and function. Earlier atomic force microscopy (AFM)-based experiments in this direction characterized the nanoscale structural features and dynamics of both canonical and centromeric nucleosomes (13, 14) as well as their assembly on different DNA sequences (15). These studies revealed greater stability of centromeric nucleosomes under in vitro conditions, as indicated by their longer dwell times and the persistence of an intact histone core after DNA dissociation (13). However, magnetic-tweezer experiments revealed broadly similar force-dependent disassembly behavior for CENP-A and H3 nucleosomes on both random and human centromeric *α*-satellite DNA, although CENP-A exhibited a modest increase in nucleosome lifetime on centromeric DNA (16).

The majority of experimental studies have focused on strongly positioned nucleosomes assembled on selected high-affinity DNA templates, such as Widom 601 sequence, possessing exceptionally high affinity for the histone octamer. Because the underlying DNA sequence determines intrinsic DNA mechanics, including its bending and deformability (17–19), it can substantially influence nucleosomal DNA conformation, DNA-histone interactions, and nucleosome dynamics(20, 21) in comparison to *α*-satellite DNA sequences. For example, *Strömberg et al.* (15) reported that CENP-A nucleosomes assembled on *α*-satellite DNA exhibit distinct positioning and wrapping dynamics compared to nucleosomes assembled on non-specific DNA. In addition, *Winogradoff et al.* (10) demonstrated that CENP-A-containing nucleosomes exhibit greater local and global fluctuations and increased DNA instability near the pseudo-dyad compared with their canonical H3-containing counterparts. Recent cryo-EM studies by *Zhou et al.* (22) showed that CENP-A nucleosomes reconstituted with Widom 601 and ASP DNA adopt similar overall nucleosome architectures, despite sequence-dependent differences in the dynamics of the terminal 10 bp of DNA. Moreover, these sequence differences were indicated to not substantially alter CENP-N-mediated nucleosome stacking. The sequence-dependent dynamics of canonical nucleosomes is known in the field. *Khatua et al.* (23) performed 12 *μ*s all-atom MD simulations and reported sequence-dependent dynamical heterogeneity between the ASP and Widom 601 sequence. The ASP nucleosome exhibited loop formation around superhelical rotation (SHL) 5 whereas the Widom 601 nucleosome showed pronounced DNA-end breathing motion at SHL 7. De Pablo’s group showed that nucleosome reposition can occur via by loop propagation or twist diffusion using "3 Site Per Nucleotide" (3SPN) coarse-grained (CG) model of DNA (24) and such mechanisms depend on DNA sequence. A recent study (25) employing enhanced-sampling MD simulations at both atomistic and CG resolutions demonstrated that in canonical nucleosomes the nucleosomal DNA unwrapping pathways are strongly sequence-dependent. Comparable studies focusing on the sequence-dependent dynamics of centromeric nucleosomes and differences between canonical and centromeric nucleosomes remain scarce. However, the studies mentioned suggest that both DNA sequence and histone identity influence nucleosome dynamics. The systematic comparison of molecular bases shaping nucleosome dynamics and how this dynamics influence the histone-DNA interactions, as well as centromeric chromatin folding, is not fully understood.

Experimental approaches remain limited in their ability to simultaneously resolve atomistic histone-DNA interactions, transient conformational fluctuations, and sequence-dependent dynamical heterogeneity across biologically relevant timescales. Detailed all-atom simulations are computationally demanding and often exclude histone tails (10), which are intrinsically disordered, but play critical roles in regulating nucleosome stability, DNA accessibility, and chromatin dynamics (26). These limitations motivate the use of CG simulation approaches for studying nucleosomes (27–31). Such approaches enable the exploration of nucleosome dynamics over longer timescales and larger spatial scales while retaining essential biophysical features of nucleosome organization.

In the present study, we employ molecular dynamics (MD) simulations with the SIRAH (Southamerican Initiative for a Rapid and Accurate Hamiltonian) (32, 33) CG force field to compare the nanoscale structural features of centromeric and canonical nucleosomes by analyzing histone core stability, histone tail-DNA interactions, and DNA structural changes. Both nucleosome types are examined using the ASP and Widom 601 DNA sequences. We expect DNA sequence to primarily influence nucleosomal DNA dynamics and histone-DNA interactions, whereas histone variant identity to play a dominant role in determining histone core structural stability. We further investigate the impact of the centromeric partner protein CENP-N on the structural dynamics and histone-DNA interactions of centromeric nucleosomes assembled on both DNA sequences. Our simulations reveal that centromeric nucleosomes exhibit reduced histone core structural plasticity compared with canonical H3 nucleosomes, while DNA sequence differentially modulates histone tail dynamics and their interactions with nucleosomal DNA. Overall, this study provides a comparative framework for understanding how DNA sequence, histone variant identity, and CENP-N binding influence nucleosome structural dynamics over microsecond timescales, offering molecular-level insights into their potential contributions to centromeric chromatin organization.

## METHODS

### System preparation and simulation protocol

The cryo-EM structure of the CENP-A containing nucleosome (22) (RCSB PDB ID: 7U46) and the crystal structure of the canonical H3 nucleosome (34) (RCSB PDB ID: 1KX5), both containing the same 147-bp palindromic *α*-satellite-derived nucleosomal DNA sequence (ASP), were used as starting models for the simulation. The missing histone tails in the centromeric nucleosome were modeled using AlphaFold3 (35). For the systematic comparison between different systems, we used previously generated trajectories of canonical nucleosomes assembled on the ASP and Widom 601 sequences (36), as well as the CENP-A nucleosome assembled on the ASP sequence (37). To investigate the effect of CENP-N binding on nucleosomal DNA, we also analyzed previously generated trajectories of the ASP-containing CENP-A nucleosome in complex with CENP-N (37) in addition to new simulations performed using the Widom 601 sequence.

Although the canonical nucleosome with Widom 601 sequence has been simulated in our earlier study (36), we have not yet explored the corresponding centromeric nucleosome. Thus, in the present work, we performed simulations following previous computational procedure (36, 37) and using the PDB ID structure 3LZ0 (38). To ensure that this modeling does not introduce structural bias, we further compared the modeled system with the experimentally determined cryo-EM structure of the centromeric nucleosome with Widom 601 DNA (PDB ID: 6TEM) (39). The details of the simulations are provided in the Supplementary Information (SI). In short, firstly, all atomistic structures were converted into their corresponding CG representations using the SIRAH Tools (40) mapping procedure. The resulting systems were then solvated with the WT4 water model (41) and maintained to a physiological concentration of 0.15 M NaCl. System specific details such as number of CG beads, solvent and ions are given in the SI (Table S1). Following energy minimization and equilibration, three independent production simulations were performed for each system using a 20 fs time step under periodic boundary conditions. Each centromeric system was simulated for 10 *μ*s, whereas each canonical system was simulated for 6 *μ*s. For consistent comparison, the final 5 *μ*s of each trajectory were used for analysis. The summary of the simulated nucleosome systems is provided in Table 1. Further, atomistic reconstructions were obtained through backmapping using the SIRAH-Backmap toolkit (40). These structures were subsequently energy minimized using the ff14SB force field within AmberTools. A detailed description of the CG preparation workflow, equilibration strategy, simulation setup, and backmapping protocol is provided in the SI.

**Table 1:** Summary of the nucleosome systems simulated and analyzed in this study. Previously reported simulation trajectories are indicated by the corresponding citations.

| **System** | **Centromeric** |  | **Canonical** |  |
|---|---|---|---|---|
|  | **ASP** | **Widom 601** | **ASP** | **Widom 601** |
| Nucleosome | 3 replicas × 10 *µ*s (37) | 3 replicas × 10 *µ*s | 3 replicas × 6 *µ*s (36) | 3 replicas × 6 *µ*s (36) |
| Nucleosome with CENP-N | 3 replicas × 10 *µ*s (37) | 3 replicas × 10 *µ*s | - | - |

### Analysis of nucleosome dynamics

To understand structural changes in the nucleosome, we employed a set of analyses described here. Global stability of the protein and DNA was monitored through the root-mean-square deviation (RMSD) of the structure with respect to the energy-minimized structure. Local flexibility was captured using root-mean-square fluctuations (RMSF). We considered backbone beads for the analysis. The compactness of histone and DNA was quantified using the radius of gyration (*R_g_*), defined as the mass-weighted root-mean-square distance of all beads from the center of mass.

The sequence-specific interactions between histone tails and DNA were characterized by calculating tail-DNA contacts (*N_C_*). Unless otherwise specified, the analysis was performed on back-mapped all-atom (AA) structures reconstructed from the trajectories obtained using coarse-grained simulations. A contact was defined when any two non-hydrogen atoms, one from the histone tail and the other from DNA, were within 4.5 Å of each other, following previous studies (42). The residue-wise DNA-contact occupancy, *P*^contact^, was calculated as the fraction of analyzed frames in which residue *i* formed at least one contact with DNA and was subsequently averaged over the three independent replicas. Thus, 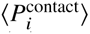 = 0.5 indicates an average DNA-contact occupancy of 50% across the independent replicas. Residues with 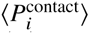 ≥ 0.5 were considered persistently DNA-contacting residues.

The time evolution of each selected parameter, 〈*X* (*t)〉*, was obtained by averaging the corresponding values across *n* = 3 independent replicas at each time point. A single time-averaged value, 〈*X* 〉, was subsequently obtained by block averaging the replica-averaged time series over the 5 *μ*s of the simulations. All analyses were performed using GROMACS tools, while in-house scripts were employed to calculate histone tail-DNA contacts. All analyses are performed using final 5 *μ*s of the trajectories based on the convergence of RMSD and *R_g_* of both histone core and DNA (see Fig. S1 and Fig. S2).

To further characterize DNA structural changes and elucidate sequence-dependent conformational heterogeneity, we calculated several established base-pair and base-pair step parameters from the backmapped atomistic trajectories. These included: (i) major and minor groove widths, and (ii) helical base-pair step (inter base-pair) parameters, which can be classified into translational parameters: shift, slide, and rise and rotational parameters: tilt, roll, and twist. Together, these parameters provide a quantitative description of local DNA geometry, flexibility, and sequence-dependent deformations within the nucleosome. All analyses were performed using the Curves+ software (43) on AA backmapped structures. In addition, DNA backbone traces were generated for each trajectory frame after least-squares alignment with the initial reference structure using the DNA backbone phosphorus atoms. The aligned backbone coordinates were projected onto the x-y plane, and the traces from all MD frames were overlaid with the initial structure to visualize DNA backbone motion and conformational fluctuations.

Errors were estimated for all quantities where applicable and are reported as the standard error of the mean (SEM), calculated as 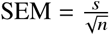, where *s* is the standard deviation of the mean values obtained from the individual replicas and *n* is the number of replicas (*n* = 3).

## RESULTS AND DISCUSSION

### Nucleosomes wrapped on the ASP DNA sequence

In the present study, we first examine the structural changes of nucleosomal DNA in centromeric and canonical nucleosomes based on the ASP sequence, followed by an assessment of histone core dynamics and histone tail interactions with ASP DNA. We then investigate how CENP-N binding modulates changes of the nucleosomal DNA and histone-DNA contacts in case of the centromeric nucleosome. Later, we evaluate the extent to which the observed dynamic features are influenced by the change of the DNA sequence from ASP to Widom 601. Unless otherwise stated, the centromeric nucleosome is represented in green, whereas the canonical nucleosome is shown in orange. Fig. 1a shows a schematic representation of both nucleosomes, highlighting the shortened *α*N helix of CENP-A. Consistent with their overall structural similarity, superposition of the corresponding crystal structures reveals only minor conformational differences, with a backbone RMSD of approximately 2 Å. Despite sharing a common nucleosomal architecture, the replacement of canonical H3 with the centromere-specific histone variant CENP-A is expected to alter the structural and dynamical landscape of the nucleosome. To understand region specific analysis of nucleosomal DNA, the DNA was described using superhelical locations (SHLs), extending from SHL 7 to 7 around the histone core (see Fig. 1b). The corresponding 147 bp DNA sequence was further mapped onto these SHLs, with base pair 74 positioned at the nucleosomal dyad (SHL 0) and the two complementary strands represented in antiparallel orientation (see Fig. 1c).

**Figure 1:**
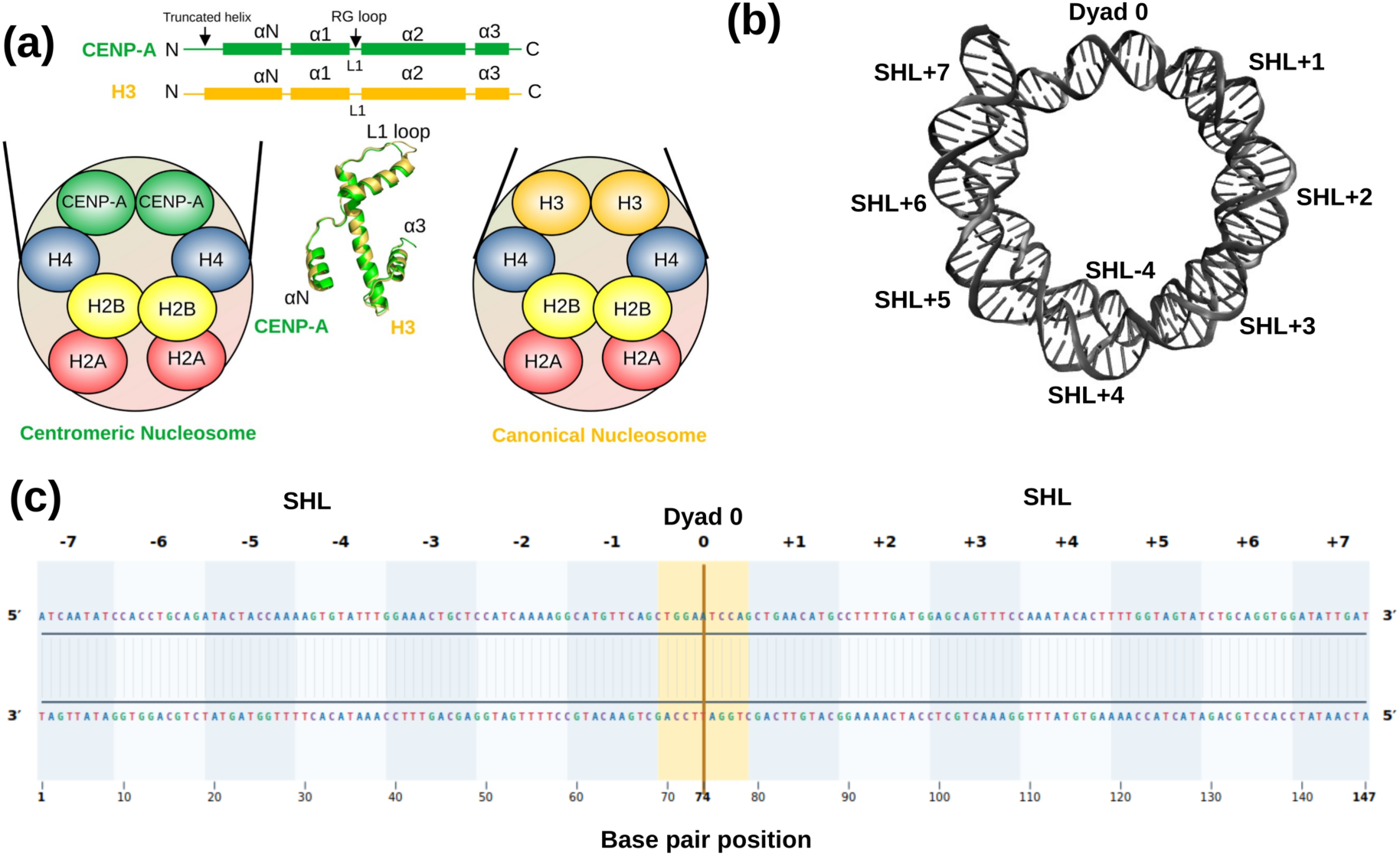
Schematic representation of (a) the canonical H3 nucleosome containing two copies of H3, H4, H2A, and H2B histones with 147 bp of DNA wrapped around the histone core, and the CENP-A centromeric nucleosome, in which CENP-A replaces H3 while retaining two copies of H4, H2A, and H2B. (b) Structural representation of the superhelical locations (SHLs) along nucleosomal ASP DNA sequence. (c) Nucleotide level mapping of the 147 bp DNA sequence onto SHLs 7 to 7. The two complementary DNA strands are shown, with base pair 74 defining the nucleosomal dyad at SHL 0.

### Global conformational dynamics of the nucleosomal DNA

To assess structural and dynamical differences between centromeric and canonical nucleosomes, we first examined global structural properties of the nucleosomal DNA with the ASP sequence. Since the same ASP sequence is used in both systems and often the centromeric nucleosomes are reconstructed in cryo-EM using the canonical templates (11, 22), the overall wrapping geometry is expected to be largely conserved. Consistent with this, the major and minor groove widths do not show significant differences between the two nucleosomes, as shown in Fig. 2a, which presents the corresponding values in bar representation. However, the canonical nucleosome DNA exhibits a slightly higher *R_g_*, indicating a marginal increase in DNA expansion while preserving the overall wrapping topology (Fig. 1b, Fig. 2c-d). The time evolution of these structural properties is provided in the SI (Fig. S2 and Fig. S3). The RMSD profiles for both systems are depicted Fig. S1 c-d, while the corresponding average values are shown in Fig. 2b in a bar representation. Overall, no substantial differences are observed, as the RMSD primarily reflects the global fluctuations of DNA.

**Figure 2:**
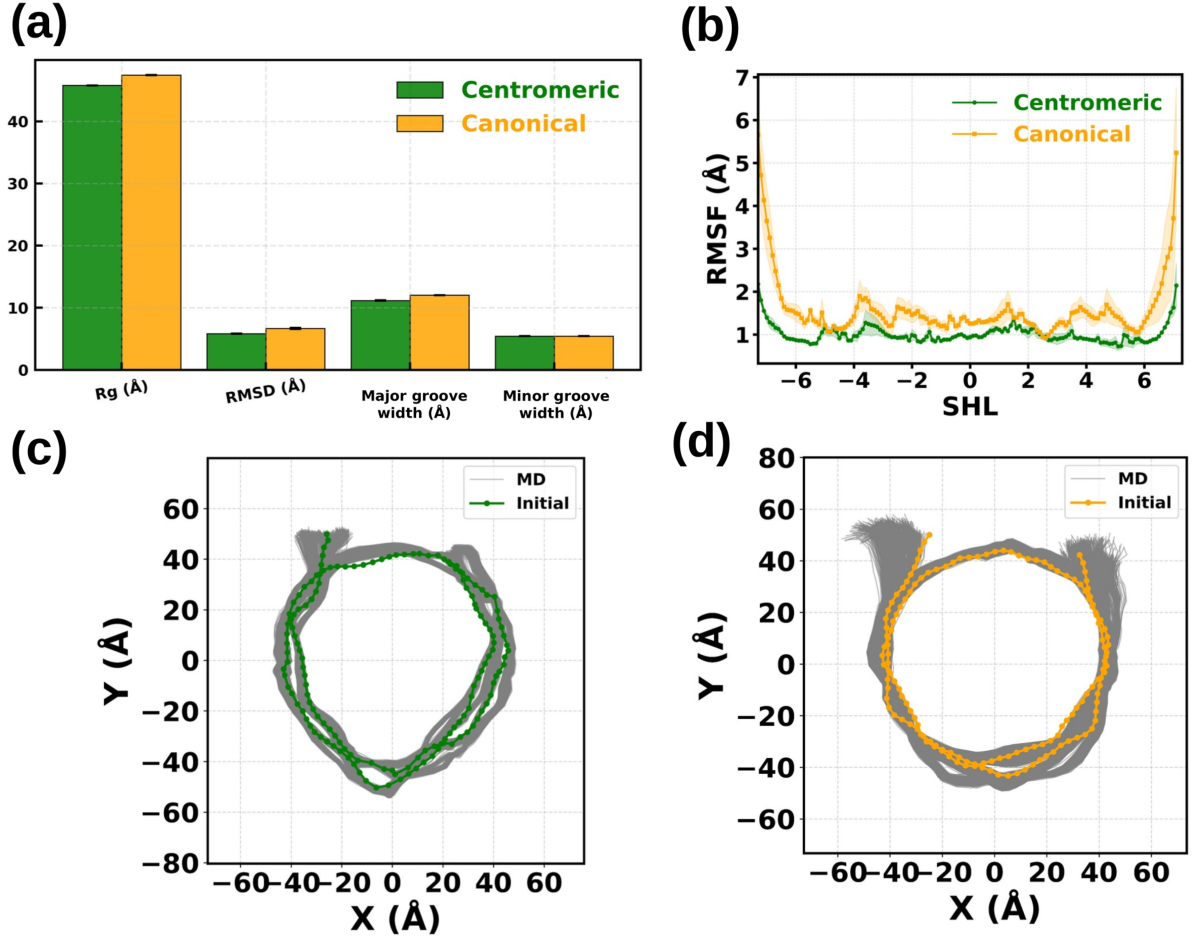
(a) Comparison of the DNA structural properties: radius of gyration (Rg), RMSD, major groove width, and minor groove width in centromeric and canonical nucleosomes. (b) RMSF of DNA as a function of base pair position mapped onto SHL. Backbone trace of nucleosomal DNA for (c) centromeric and (d) canonical nucleosome. The dotted line indicates initial structure while DNA conformations obtained from independent simulations are marked in grey.

To obtain a more detailed picture of local DNA flexibility, RMSF of individual base pairs as a function of their superhelical location (SHL) was calculated. Figure 2b shows the average RMSF of DNA in the centromeric and canonical nucleosomes, represented in green and orange, respectively. The shaded regions indicate statistical errors obtained from three replicas. In general, the canonical nucleosome exhibits a slightly higher RMSF values than the centromeric nucleosome, however, the differences are rather small considering the CG level of the molecular representation. The differences are particularly pronounced at the DNA entry-exit regions, where distinct DNA wrapping and terminal flexibility have been shown to differ between canonical and centromeric nucleosomes (44). The two-dimensional DNA backbone traces further reveal a broader distribution of terminal DNA conformations for the canonical nucleosome, indicating greater conformational fluctuations of the DNA ends under the simulated conditions (Fig. 2c,d). However, the enhanced conformational variability can be influenced by one canonical nucleosome replica exhibiting somewhat larger structural deviations (see Fig. S1). Thus, the differences may be related to the sampling of terminal DNA conformational states rather than an intrinsic difference in DNA-end flexibility between the two nucleosome types.

### Structural plasticity of the histone core

Earlier studies have suggested that the structural dynamics of the histone octamer are closely coupled to nucleosomal DNA unwrapping and breathing (42, 46, 47). In particular, restricting the conformational flexibility of the histone octamer core has been shown to suppress DNA fluctuations and reduce the extent of DNA unwrapping (42, 48, 49). Therefore, characterizing the fluctuations of the histone core is crucial for understanding the relationship between histone dynamics and nucleosome stability in both centromeric and canonical nucleosomes. Since histone tails are intrinsically disordered and exhibit large-amplitude fluctuations, the present analysis focuses exclusively on the histone core domains. This approach provides an assessment of the intrinsic conformational plasticity of the nucleosome and its potential role in modulating nucleosome stability and DNA dynamics.

Figure 3a-d shows the residue-wise RMSF of the backbone CG beads for one copy (chain 1) of each histone core protein in the centromeric and canonical nucleosomes. Only chain 1 is presented in the main text for clarity, as the two histone copies exhibit broadly similar RMSF profiles, except for canonical H3. The corresponding profiles for chain 2 are provided in the SI (Fig. S4). From Fig. 3a, it is seen that CENP-A exhibits lower RMSF compared to the canonical H3 the main core region. However, the L1 loop region (Fig. 1a) exhibits elevated fluctuations in both nucleosomes. In each system, this region is more flexible than the adjacent *α*-helical segments, consistent with its loop architecture. A more pronounced difference is observed in the *α*N helix near the N-terminal tail. In chain 1, the canonical H3 *α*N helix exhibits RMSF values more than 1 Å higher than the corresponding region of CENP-A, whereas this difference exceeds 2 Å in chain 2 (Fig. S4). By contrast, the *α*1 region (Fig. 1a) shows only minor differences (less than 1 Å) between the two nucleosomes. The chain-dependent increase in *α*N flexibility may reflect asymmetric interactions between the H3 N-terminal tails and their neighboring DNA ends. Previous simulations have shown that asymmetric nucleosomal DNA breathing is accompanied by distinct histone tail-DNA interactions at the two entry and exit regions (50). Thus, the elevated and asymmetric *α*N fluctuations observed in the canonical nucleosome are consistent with differential coupling between the H3 *α*N/tail region and terminal DNA. In the centromeric nucleosome, our previous study showed that CENP-N binding near the CENP-A L1 loop enhances fluctuations in the N-terminal region of CENP-A (37), therefore, in next sections, we further examine histone tail-DNA interactions, as well as the impact of CENP-N.

**Figure 3:**
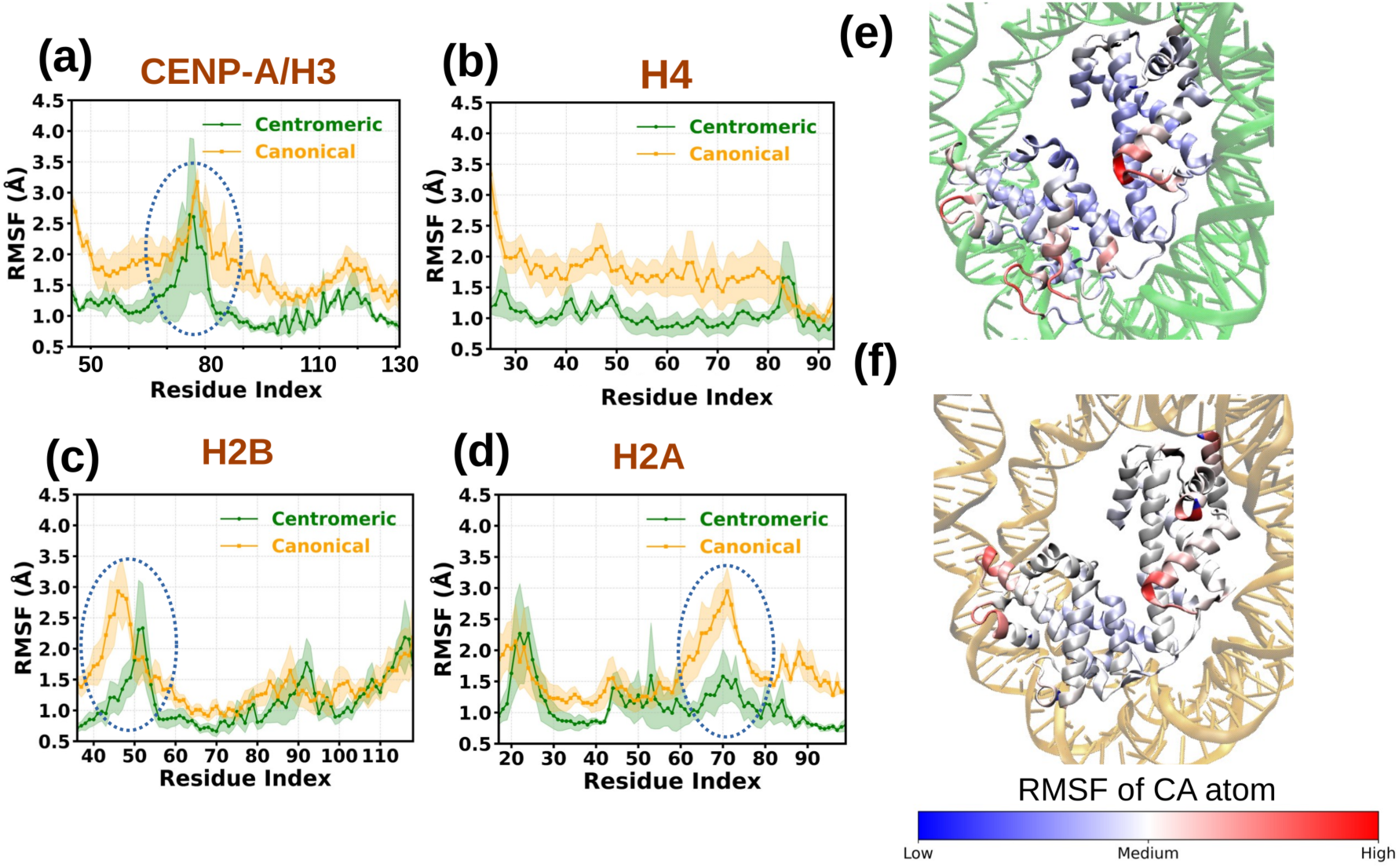
Root mean square fluctuations (RMSF) of the histone core: (a) CENP-A (centromeric) / H3 (canonical), (b) H4, (c) H2B, and (d) H2A histones. Solid lines represent the mean RMSF values averaged over three independent simulation replicas using the final 5 *μ*s of each trajectory, while the shaded regions indicate the corresponding standard error of the mean across the replicas. The residue wise fluctuations of backbone CA atom are further mapped onto the equilibrated structures of the (e) centromeric and (f) canonical nucleosomes, where residues are colored according to their relative RMSF values. Red indicates regions with higher fluctuations, whereas blue denotes more rigid, less fluctuating, regions. The RMSF based coloring was generated in VMD (45) using a user defined color scale. Results for one histone copy are presented. The corresponding data for the second copy is depicted in Fig. S4.

While CENP-A/H3 is the key determinant distinguishing centromeric and canonical nucleosomes, histones H4, H2A, and H2B are conserved core components present in both systems. Nevertheless, the distinct dynamics of CENP-A and H3 can influence the dynamics of these conserved histones by altering inter-histone interactions and their coupling with nucleosomal DNA. The H4 histone RMSF profile indicates enhanced flexibility in the histone core for canonical system (see Fig. 3b). No significant differences in RMSF are observed for H2B (see Fig. 3c) and H2A (see Fig. 3d), except for certain highlighted loop regions in both proteins. However, greater differences occur for the second copy of those histones (Fig. S4). Overall, the CG-MD simulations indicate greater local conformational rigidity within the histone core of the CENP-A nucleosome than in its canonical H3 counterpart. This finding is consistent with hydrogen-deuterium exchange measurements by *Black et al.* (51), which specifically revealed increased conformational constraints in regions forming the CENP-A-H4 interface. Furthermore, to demonstrate the differences in the global structural plasticity of the core histone, the residues of copy 1 histones were mapped onto the equilibrated structures according to their RMSF values, which is visualized in Fig. 3e and f for the centromeric and canonical nucleosomes, respectively. Lower structural plasticity in the centromeric nucleosome is clearly seen by a higher extend of the lower RMSF values presented in blue.

### Histone tails-DNA interactions

Histone tails play a crucial role in regulating nucleosome structure and dynamics through their extensive interactions with DNA and may contribute to internucleosomal interactions, thereby influencing higher-order chromatin structure (26, 52–55). Owing to their intrinsic flexibility, the tails can establish both persistent and transient contacts with the DNA backbone, thereby influencing processes such as DNA breathing, unwrapping, and stacking (23, 52, 56–58). Given the differences in DNA flexibility and histone core dynamics observed between the canonical and centromeric nucleosomes, we next examined the contribution of histone tails by quantifying the contacts between individual histone tails and nucleosomal DNA for both systems. For that, the average number of contacts, 〈*N_C_*〉 (see Methods), was calculated. The time evolution of 〈*N_C_* (*t)*〉, obtained by averaging over multiple replicas at each time point, are shown in the SI (Fig. S5 a-e) together with the corresponding error estimates. The trajectory-averaged contact numbers obtained using block averaging are listed in Table 2. Fig. 4a-b shows representative structural conformations of the CENP-A/H3 N-terminal tail in both systems sampled in MD.

**Figure 4:**
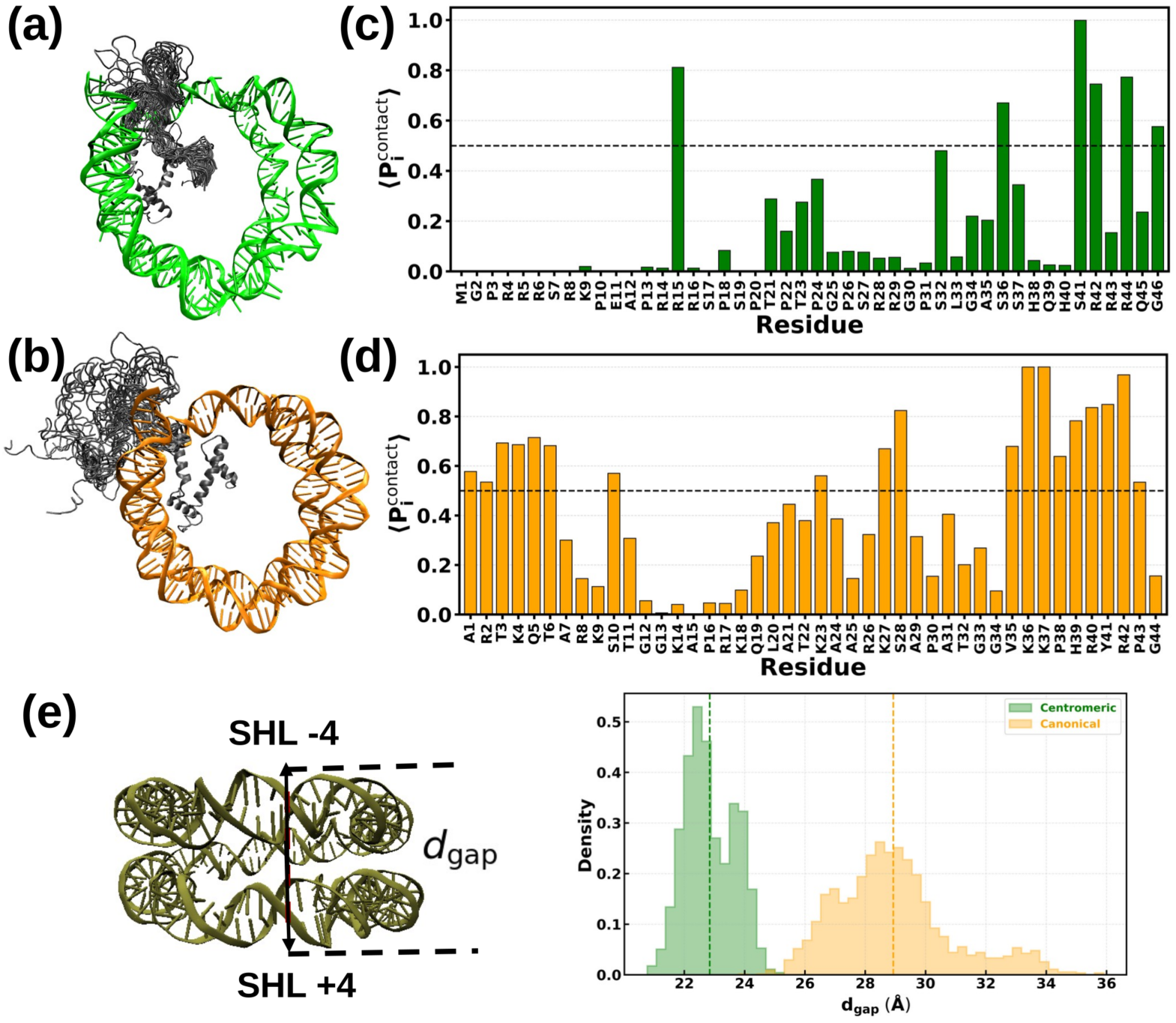
Representative structural conformations of the (a) CENP-A and (b) H3 N-terminal tails obtained from three independent simulation runs. Residue-wise average DNA contact occupancy plot for (c) CENP-A and (d) H3 histone N-terminal tails. Averaging was performed over multiple replicas. (e) The illustration of the gaping distance (*d_gap_*) in the nucleosome together with the *d_gap_* distribution for centromeric and canonical nucleosomes obtained in CG-MD simulations.

**Table 2:** Average histone tail-DNA contacts ( *N_C_*) for centromeric and canonical nucleosomes assembled with ASP and Widom 601 DNA. Values in parentheses represent the minimum–maximum range obtained from the corresponding time-evolution profiles. For the Widom 601 systems, ↑ and ↓ indicate an increase and decrease, respectively, relative to the corresponding ASP system, whereas ∼ indicates a change of less than 10%.

| Histone type | ASP sequence |  | Widom 601 sequence |  |
|---|---|---|---|---|
|  | Centromeric | Canonical | Centromeric | Canonical |
| CENP-A/H3 | 80.79 (*55–102*) | 345.08 (*298–410*) | ↑ 188.56 (*171–225*) | ↓ 281.66 (*220–321*) |
| H4 | 128.04 (*70–153*) | 192.17 (*159–230*) | ∼ 132.44 (*112–152*) | ↓ 152.15 (*126–173*) |
| H2B | 303.13 (*270–325*) | 352.33 (*315–385*) | ↓ 216.41 (*197–247*) | ∼ 329.02 (*299–382*) |
| H2A | 201.14 (*174–249*) | 156.95 (*133–215*) | ∼ 190.56 (*172–226*) | ∼ 166.32 (*125–182*) |
| H2A C-terminal | 60.13 (*44–81*) | 44.47 (*25–61*) | ↑ 129.59 (*101–151*) | – |

From Fig. 4a-b, a significant difference between the behavior of the histone tails in centromeric and canonical nucleosomes are visible, with the canonical system exhibiting a sufficient variety of interaction poses and contacts with the DNA. This explains a much higher number of contacts between the tails and the DNA compared to the centromeric system, where on average 345 histone-DNA contacts were detected in comparison to only 80 contacts between the CENP-A tails and the DNA (see Table 2). In the centromeric system, the CENP-A N-terminal tail is only partially engaged with DNA, whereas in the canonical nucleosome, the H3 N-terminal tail directly interacts with the DNA entry/exit region. This observation is consistent with previous simulation studies, where the H3 N-terminal tail was found to promote DNA unwrapping through direct DNA interactions (23), whereas the CENP-A N-terminal tail predominantly forms more persistent contacts with the H2A histone (37). To further characterize these differences at the residue level, we calculated the average DNA-contact occupancy, 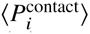, from the backmapped trajectories. The CENP-A tail (see Fig. 4c) showed more localized DNA association, with only 6 of 46 residues exhibiting 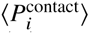 ≥ 0.5, compared with 19 of 44 residues for the H3 tail (Fig. 4d). The difference was particularly pronounced at the extreme N terminus: residues A1 to T6 of H3 exhibited persistent DNA contacts, whereas the first 14 residues of CENP-A (M1 to R14) remained essentially disengaged from DNA. This behavior suggests distinct interaction preferences of the two histone tails within the nucleosome. CENP-A and DNA contacts were instead localized further along the tail, with persistent contacts involving R15 and residues within the S36 to G46 region. Thus, the residue wise analysis supports the structural observation that the canonical H3 tail establishes more extensive interactions with DNA, whereas DNA contacts of the CENP-A tail are comparatively localized.

The N-terminal tails of H4 (around 23 amino acid long) and H2B show a similar trend (see Table 2), although the differences between the centromeric and canonical systems are less pronounced than those observed for CENP-A/H3. For H4, the centromeric nucleosome exhibits an average of approximately 128 tail-DNA contacts, compared with approximately 192 contacts in the canonical nucleosome. Residue wise contact occupancies nevertheless show that H4 remains substantially engaged with DNA in the centromeric system, with 11 of 24 analyzed residues exhibiting a 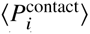 value of at least 0.5 (see Fig. S6a). 12 out of 24 amino acids in the N-terminal exceeds this threshold in the canonical system (see Fig. S6b). Thus, the lower 〈*N_C_*〉 in the centromeric nucleosome does not reflect a complete disengagement of the H4 tail from DNA, but rather differences in the extent and distribution of H4-DNA contacts.

A similar behavior is observed for H2B N-terminal tail. In the centromeric nucleosome, H2B exhibits an average of approximately 303 tail-DNA contacts, compared with approximately 352 contacts in the canonical system. Consistent with this relatively modest difference, extensive H2B-DNA association is retained in both systems, with 15 of 37 analyzed residues in the centromeric nucleosome exhibiting 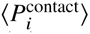 ≥ 0.5, compared with 16 of 35 residues in the canonical nucleosome (see Fig. S6 c-d). Several H2B residues in both systems exhibit 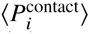 values approaching unity, indicating highly persistent DNA association. Thus, unlike the pronounced difference observed between the CENP-A and H3 N-terminal tails, the H4 and H2B tails remain extensively associated with DNA in both nucleosome types, while the lower 〈*N_C_*〉 in the centromeric system reflects differences in the extent and distribution of these tail-DNA contacts rather than their overall loss.

For histone H2A, we further report the average contacts for both the N- and C-terminal tails, which are approximately 16 and 12 amino acids long, respectively, as both contribute to nucleosome dynamics. In contrast to the H4 and H2B tails, H2A exhibits a higher number of tail-DNA contacts in the centromeric nucleosome for both terminal regions (see Fig. S7). The difference is particularly evident for the N-terminal tail, which exhibits an average of approximately 201 DNA contacts in the centromeric nucleosome compared with approximately 156 contacts in the canonical system. Consistent with this increase, the residue-wise analysis reveals more persistent DNA association in the centromeric system (see Fig. S7a-b). The H2A C-terminal tail also shows a higher average number of DNA contacts in the centromeric nucleosome than in the canonical system (Table 2). However, this increase is not accompanied by a larger number of persistently DNA-contacting residues as depicted in Fig. S7c-d.

Overall, the results obtained reveal a redistribution of histone tail-DNA interactions between the canonical and centromeric nucleosomes, rather than a uniform change across all histone tails. While CENP-A/H3 exhibits the most pronounced difference, with substantially more extensive H3-DNA interactions in the canonical nucleosome, H2A shows the opposite trend, with enhanced N- and C-terminal DNA contacts in the centromeric system. To validate the consequence of the latter, we further analyzed DNA gaping motion, which is distinct from the nucleosome breathing (i.e., changes in the separation between the two DNA gyres). Previous simulation studies (42) have shown that the dynamics of the H2A N-terminal tail are coupled to DNA gaping motion. This conformational transition was first observed experimentally by *Ngo et al.* (59) using single-molecule FRET and was proposed theoretically earlier by *Mozziconacci et al.* (60) as a mechanism to facilitate the tight packing of higher-order chromatin structures. During this transition, the two half-turns of nucleosomal DNA (SHL+ and SHL-) move apart along an axis perpendicular to the nucleosome plane. In the present study, we quantified the gaping motion using the gaping distance (*d_gap_*), defined as the distance between the DNA segments located in the SHL-4 and SHL+4 regions (see Fig. 4e). Across multiple replicas, it is higher and broader for the canonical nucleosome than for the centromeric nucleosome (see the average distribution of 〈*d*_gap_〉 on the right in Fig. 4e). The time evolution of 〈*d_gap_* (*t*)〉 is depicted in SI in Fig. S5 f. Significant difference is observed between the systems studied with canonical nucleosome being much more dynamic: the 〈*d_gap_* (*t)*〉 values between 27 and 31 Å and with rather high fluctuations are present. The centromeric system follows more stable motions with the most populated *d_gap_ t* values of around 22 Å. This observation follows trends of the H2A N-terminal tail contacts with the DNA, agreeing well with the observations reported in the literature(42), suggesting a potential role of these interactions in modulating DNA gaping dynamics. Notably, the smaller DNA gaping distance in the centromeric system is consistent with the reduced DNA and histone-core fluctuations observed in the RMSF analysis. Together, these observations indicate more closely spaced DNA gyres with a more restricted deformation and reduced local conformational fluctuations in the simulated CENP-A. This reduced DNA gaping and local fluctuations may contribute to maintaining specific CENP-A nucleosome structural integrity, potentially supporting the persistence of CENP-A nucleosomes as platforms for centromere-associated protein assembly. However, their consequences for protein recruitment and resistance to mechanical disruption cannot be established from these structural analyses alone.

### Impact of CENP-N binding on the nucleosomal DNA

A key function of the centromeric nucleosome is to serve as a platform for the assembly of inner kinetochore proteins through a network of protein-protein interactions (12, 61). Among these, CENP-N acts as a specific reader of CENP-A-containing nucleosomes and plays an important role in centromere recognition (12, 62). Its impact on reshaping conformational states of CENP-A we reported previously (37). However, CENP-N interacts with both the CENP-A histone and specific DNA base pairs located between SHL+2 and SHL+3.5 (Fig. 1c) as depicted in Fig. 5a. Thus, here we focus on the DNA-related structural differences and dynamical changes in the binding interface. Firstly, we calculated the contacts between DNA and the CENP-A histone within the SHL+2 to SHL+3.5 region (see in green in Fig. 5b) and later we compared it to the H3 histone-DNA contacts in the canonical nucleosome, as well as to the contacts in the functional state of the CENP-A nucleosome, i.e., upon the complexation with CENP-N.

**Figure 5:**
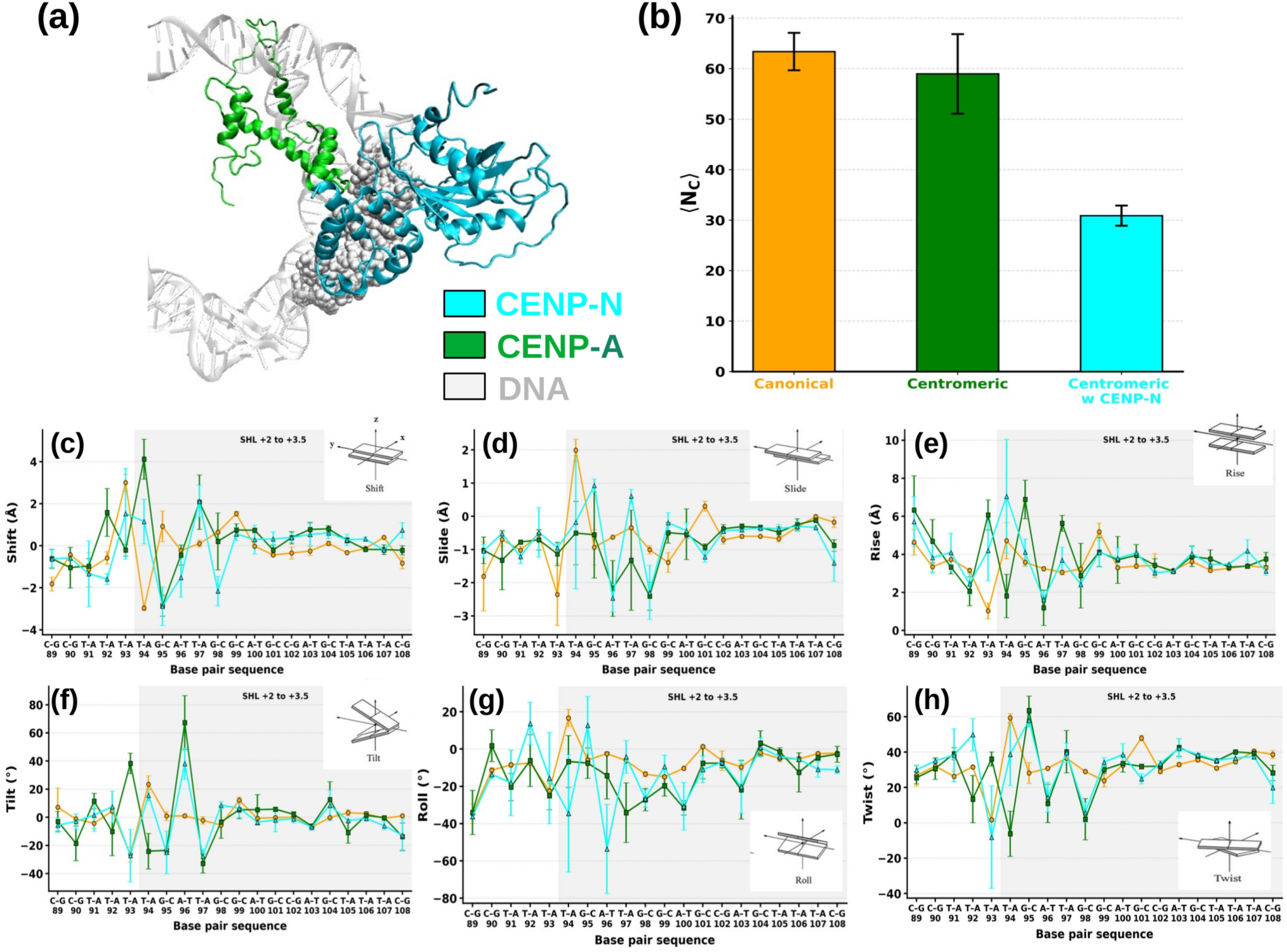
(a) Binding interface between the centromeric nucleosome and the kinetochore protein CENP-N. CENP-N is shown in cyan, while the DNA is in gray. The DNA region involved in CENP-N binding (SHL 2–3.5) is highlighted using a van der Waals (vdW) representation in VMD. (b) Bar plots showing the average number of contacts ( *N_c_*) between the SHL 2-3.5 DNA region and the H3 (canonical) or CENP-A (centromeric) histone core. Changes in the DNA inter-base-pair parameters: Shift (c), Slide (d), Rise (e), Tilt (f), Roll (g), and Twist (h). A schematic illustration of each parameter is included in the corresponding panel. The SHL +2 to +3.5 region is highlighted in gray. The centromeric and canonical nucleosome systems are represented in green and orange, respectively, while the centromeric nucleosome bound to CENP-N is shown in cyan.

As depicted in Fig. 5b, in the canonical nucleosome an average of 63 H3-DNA contacts were detected, which is rather similar to the CENP-A-DNA contacts in the centromeric nucleosome without CENP-N. However, upon CENP-N binding, contacts between CENP-A and DNA significantly decrease by approximately 48%. The time evolution of 〈*N_C_* (*t)*〉 is shown in Fig. S8a. To identify residues associated with this decrease, we calculated the residue-wise DNA contact occupancy, 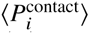, for CENP-A residues in the region from F78 to A90 (see Fig. S8b-c). Contacts formed by the residues from V82 to Q89 are the most relevant and W86, Q87 and A88 show the largest decrease in the contact occupancy after the CENP-N binding, while N85 retains high DNA contact occupancy in both systems. Thus, the average contact decrease visible in Fig. 5b can be assigned to the changes within those residues. Our previous study identified R80 and G81 in the CENP-A L1/RG loop as a major recognition site of CENP-N, which additionally interacts with nucleosomal DNA in the same SHL region (37). Neither R80 nor G81 forms persistent contacts with DNA, as revealed by the present analysis, suggesting that CENP-N recognition of the RG loop does not directly compete with DNA interactions involving these. Instead, the reduced CENP-A-DNA contact occupancy suggests a local reorganization of histone-DNA interactions, potentially mediated by conformational changes in the region adjacent to the RG loop.

To further elucidate the molecular basis of CENP-N-induced structural changes of DNA, we analyzed DNA base-pair-step parameters, which provide a detailed description of local DNA conformational dynamics. They have been widely used to characterize nucleosome structure and structural flexibility (23, 36, 63). We calculated the base-pair step parameters for the canonical, centromeric, and CENP-N-bound centromeric nucleosomes, focusing on the SHL+2 to SHL+3.5 region together with several preceding base-pair steps to assess whether the structural perturbations extend outside this region (see Fig. 1b,c). The canonical and centromeric nucleosomes exhibit largely similar DNA conformations, although differences are observed at individual base-pair steps. Shift differs mainly at the T-A (94) and G-C (95) steps belonging to the SHL+2 to SHL+3.5 region, while differences are also observed at the preceding T-A (92) and T-A (93) steps. Changes in slide extend over a broader sequence range, from T-A (93) to G-C (101), with the exception of A-T (100), whereas rise shows more localized differences. Overall, the largest differences between the canonical and centromeric nucleosomes are concentrated across the T-A (93), T-A (94), G-C (95), A-T (96), T-A (97), and G-C (98) sequence segment, approximately corresponding to the SHL +2 to +2.5 region. The rotational parameters tilt, roll, and twist (Fig. 5f-h) also show pronounced differences within this initial segment. Since the ASP DNA sequence is identical in the two systems, the observed changes in local DNA geometry can be attributed to differences in the nucleosomal environment associated with the replacement of H3 by CENP-A. The direction and magnitude of these differences vary along the DNA sequence, indicating localized and sequence-position-dependent rather than uniform structural changes.

Upon CENP-N binding, some additional local sequence-dependent changes are observed in the DNA base-pair-step parameters in comparison to unbounded CENP-A system (Fig. 5c-h). The most pronounced differences are concentrated within the DNA segment spanning positions 92-98, where several translational and rotational parameters are affected simultaneously, with smaller changes extending to neighboring steps. For the translational parameters, rise shows the largest changes across this segment except in T-A (92) and A-T (96) base pair (see Fig. 5e), while shift changes most strongly at the first T-A (94) and the following G-C (98) step (see 5c). Slide exhibits more localized changes, particularly at the G-C (95) and T-A (97) steps (see Fig. 5d). The rotational parameters also display pronounced changes: roll changes strongly across the T-A (94), G-C (95), A-T (96), and T-A (97) segment, while tilt shows particularly large changes at the first T-A (92), T-A (93), T-A (94) and A-T (96) steps, with smaller deviations extending further along the sequence. Twist exhibits its largest change at the first T-A (93), T-A (94) step, together with additional smaller changes at several subsequent sequence positions. Based on these observations, we see that CENP-N binding is associated with a localized reorganization of DNA geometry, involving changes in both base-pair stacking and local bending and twisting, rather than a uniform structural deformation along the DNA. Such changes may reflect altered local constraints imposed by CENP-N binding and the accompanying rearrangement of histone-DNA interactions.

### Effect of Widom 601 DNA on nucleosome dynamics

To assess the influence of DNA sequence, we further compared both nucleosomes assembled with Widom 601 DNA. For clarity, we depict the differences between both sequences in Fig. 6a, where the ASP and Widom 601 sequences were aligned relative to their nucleosomal dyads. Among the 145 nucleotide positions in the Widom 601 sequence, 34 contain bases identical to those in ASP, corresponding to a dyad-aligned positional identity of 23.4%, whereas 111 positions (76.6%) differ. Identical bases at corresponding positions are highlighted in red while rest are in black. The two sequences differ substantially across the SHL regions, providing distinct sequence contexts for investigating sequence-dependent nucleosome dynamics.

**Figure 6:**
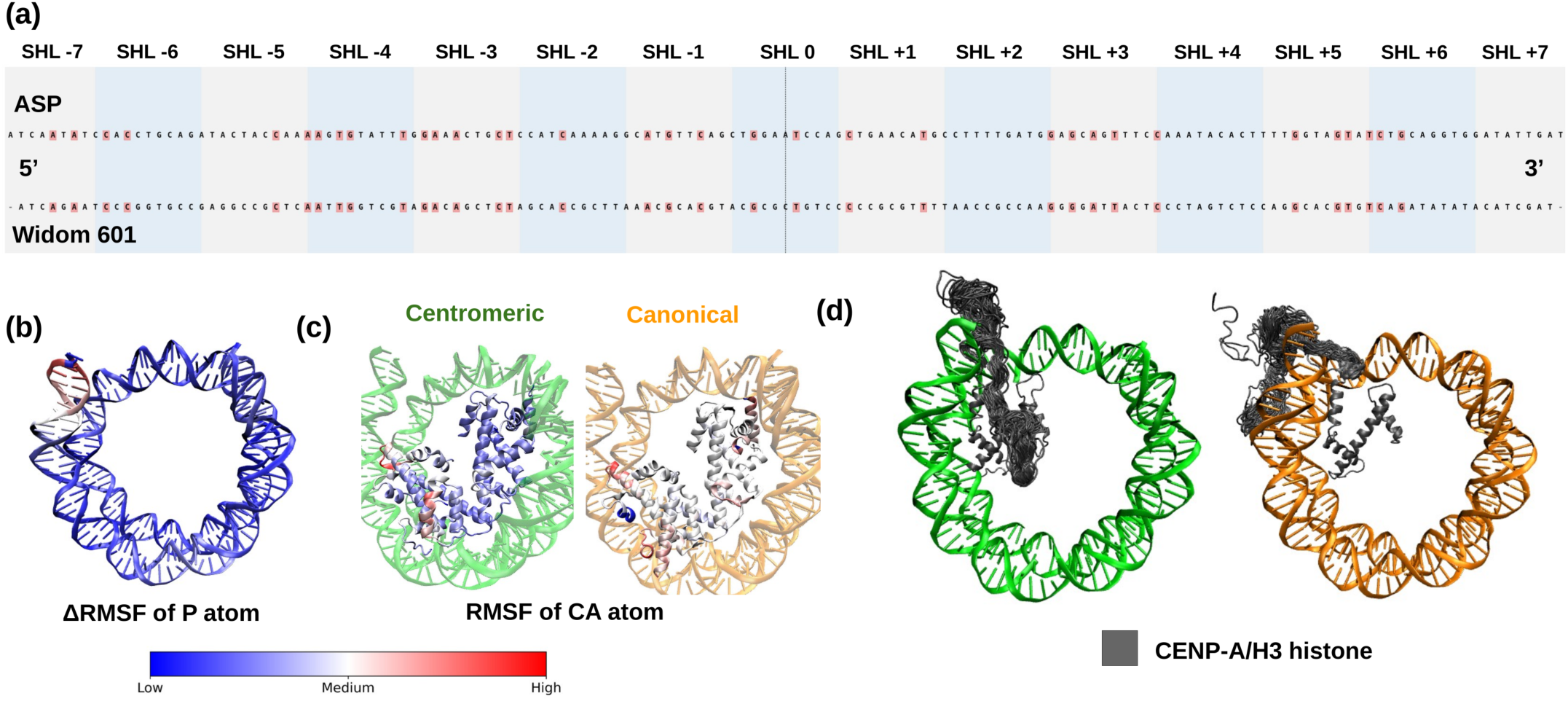
(a) Nucleotide-level mapping of the 147-bp DNA sequence onto SHLs 7 to 7 for the ASP and Widom 601 sequences. Individual nucleotide identities are shown for the 5^′^ to 3^′^ strand only. (b) Difference in RMSF (ΔRMSF) between the canonical and centromeric nucleosomes mapped over individual DNA base pairs for the Widom 601 sequence. (c) RMSF of the globular core domains of the histones. Residue-wise fluctuations are mapped onto the equilibrated structures of the centromeric (green) and canonical (orange) nucleosomes. (d) Representative structural conformations of the CENP-A and H3 N-terminal tails obtained from three independent simulation runs.

In the current study, the centromeric nucleosome model was generated by replacing the ASP DNA in the initial CENP-A nucleosome model with the Widom 601 sequence, as described in the Methods section. The experimentally resolved CENP-A nucleosome assembled with Widom 601 DNA (PDB ID: 6TEM) could not be used directly because it lacks the histone tails and does not contain the bounded CENP-N protein. To ensure that the reconstruction procedure did not introduce substantial structural artifacts, we compared the modeled system with the deposited 6TEM structure by examining histone core-DNA contacts, the DNA radius of gyration, RMSD, and residue-wise RMSF as explained in the SI (Fig. S9 and S10). These quantities showed only minor differences between the two systems, supporting the structural consistency of the reconstructed model. We subsequently used the validated model to investigate sequence-dependent effects on nucleosome dynamics.

### Structural analysis

The difference in DNA fluctuations (ΔRMSF) between the centromeric and canonical nucleosomes, mapped onto the DNA base pairs of the equilibrium structure, is shown in Fig. 6b. The differences are predominantly localized to one DNA end, whereas the nucleosomal DNA spanning SHL 5 to 5 exhibits only minimal differences in fluctuations (less than 1 Å) between the two nucleosomes (see Fig. S11). We further performed residue-wise RMSF in histone core to see whether the more stable nature of centromeric nucleosome (as observed for the ASP DNA) still preserves upon the sequence change. Fig. 6c shows the RMSF of the globular core domain of histones mapped onto the equilibrium structures for the centromeric and canonical nucleosomes. Residues are colored according to their RMSF values, as described earlier, using VMD. The centromeric system exhibits reduced fluctuations compared to the canonical system, consistent with earlier observations based on the ASP sequence (see Fig. 3e,f). Per residue RMSF for both copy of histones is further represented in SI in Fig. S12. The histone-core RMSF profiles show modest, region-specific differences between the ASP and Widom 601 systems, however the differences between the nucleosomes cores are generally smaller than observed for the nucleosomes assembled on the ASP DNA. In chain 1, both CENP-A and H3 histones exhibit an approximately 1 Å differences in RMSF with Widom 601 DNA, which is on the borderline of the coarse-grained accuracy of the used simulation method (37). Chain 2, however, shows stronger RMSF for the canonical H3 within the *α*2-helical region. For H2B, both the canonical and centromeric nucleosomes exhibit reduced RMSF values relative to their corresponding ASP systems.

The enhanced conformational plasticity of the histone core and DNA of canonical nucleosome observed in our simulations is consistent with recent experimental findings reported by Lyubchenko and co-workers (13, 64). Using time-lapse high-speed atomic force microscopy (HS-AFM), the authors compared the nanoscale dynamics of canonical H3 and centromeric CENP-A nucleosomes assembled on an identical Widom-601 DNA template. Their results demonstrated that CENP-A nucleosomes are significantly more stable than their canonical counterparts. Histone dissociation during DNA unwrapping was observed much more frequently in canonical nucleosomes, whereas CENP-A nucleosomes largely retained the integrity of the histone core during unwrapping events. In addition, CENP-A nucleosomes exhibited longer dwell times, defined as the time required for complete unraveling of the nucleosome.

### Histone tail-DNA interactions

Similarly as we performed for the nucleosomes with the ASP sequence (see Table 2), we calculated average histone tail-DNA contacts, *N_C_*, in the case of wrapping on Widom 601 DNA. The values for both systems are listed in Table 2 with the arrow signs indicating the generally observed trends between the centromeric and canonical nucleosomes with different DNA. The time evolution of histone tails-DNA contacts is depicted in Fig. S13.

Changing the DNA sequence from ASP to Widom 601 produces a marked redistribution of histone-tail–DNA contacts in both nucleosome types, however differences are histone-dependent and do not follow similar trends. In the centromeric nucleosome, the average number of CENP-A N-terminal tail contacts increases from 80.79 to 188.56 in ASP and Widom 601 DNA, respectively, indicating higher preference towards binding with Widom 601 DNA. Fig. 6d shows representative conformations of the CENP-A and H3 N-terminal tails sampled during the MD simulations. The tail dynamics exhibit trends similar to those observed in the ASP systems, with the canonical H3 nucleosome displaying a wider range of tail conformations and interaction modes with nucleosomal DNA. Although, H3 N-terminal tail–DNA contacts decrease in the presence of the Widom 601 sequence as compared to ASP which were also observed durinh unwrapping in canonical nucleosomes (25). The H2A C-terminal contacts also increase substantially, from 60.13 to 129.59, and they are significantly different between the centromeric and canonical nucleosomes (see Fig. S12c). The absence of these contacts in canonical Widom 601 system may arise from the incomplete resolution of this tail in the initial crystal structure. The H2B N-terminal contacts decrease in centromeric nucleosome with Widom 601 DNA: from 303.13 to 216.41, while differences between other systems are less pronounced. The H4 and H2A N-terminal contacts with DNA remain comparatively similar with changing the DNA sequence in both cases with centromeric nucleosomes showing generally somewhat less contacts than canonical ones, however, the differences are not so significant as seen above. Thus, from our CG-MD simulations, we clearly see that Widom 601 DNA reorganizes histone-tail-DNA contacts in both nucleosomes, introducing DNA-sequence-dependent changes in histone-DNA interactions, which are modulated by histone composition and cannot be generalized between canonical to centromeric nucleosomes.

The residue-wise DNA contact probability, 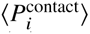, for the CENP-A N-terminal tail (see Fig. S14) indicates the largest increase of contacts originated from R16 (0.01 to 0.68) and R43 (0.15 to 1.00), with the R42-R44 region becoming persistently DNA-associated in the Widom 601 system. Similarly, the enhanced H2A C-terminal interaction is primarily associated with increased contact probabilities of S366, K371, G372, and K373, while K369 remains persistently associated with DNA in both systems. The reduced H2B N-terminal interaction is accompanied by pronounced decreases at K389 (0.99 to 0.08) and K401 (0.67 to 0.14). These residue-specific changes indicate a sequence-dependent redistribution of histone-tail interactions rather than a uniform alteration of DNA association. In contrast, for canonical nucleosome (see Fig. S15), pronounced reduction of histone-DNA contacts is observed in H3 tail for K4 (0.69 to 0.31), Q5 (0.71 to 0.18), K23 (0.56 to 0.02), Y41 (0.85 to 0.15), and R42 (0.97 to 0.35), although increased contacts at K14, R17, and K18. Similarly, the H4 K140-K147 region remains strongly DNA-associated in both systems, while the overall reduction is primarily associated with contact losses at R138, R154, and K155, partially compensated by increased contacts at K151 and R152. These results demonstrate that replacing ASP with Widom 601 DNA does not uniformly strengthen or weaken histone-tail interactions but instead redistributes the tail-DNA contact network, reducing the overall H3 and H4 tail association in the canonical nucleosome while increasing the contributions of the CENP-A N-terminal and H2A C-terminal regions in the centromeric nucleosome. These differences may arise from sequence-dependent variations in local DNA structure that alter the accessibility of individual DNA regions to the histone tails. The distinct distributions of CENP-A/H3 histone-DNA contact probabilities in the nucleosomes are consistent with recent experimental and computational evidence demonstrating histone-variant-specific contact patterns and cooperative contributions of the N-terminal tail, latch region, and *α*-N helix to nucleosomal DNA organization (65).

Finally, we examined the average DNA gap distance, 〈*d*_gap_〉, as analyzed earlier for the ASP DNA. The time evolution of 〈*d*_gap_〉 is depicted in Fig. S13f. Replacing ASP with Widom 601 DNA does not significantly impact intergyre separation of the canonical nucleosome in the region between SHL 4 to 4 captured by d-gap: the average 〈*d*_gap_〉 is 28.73 and 29.03 Å in both cases. However, the centromeric nucleosome shows a marked increase of 〈*d*_gap_〉 from 22.80 to 27.54 Å in changing the DNA sequence from ASP to Widom 601. These results indicate that DNA gaping in the canonical nucleosome is relatively insensitive to the sequence change, while the more compact DNA-gyre arrangement is observed for the centromeric nucleosome with ASP DNA, which is not maintained with Widom 601 DNA.

### Effect of CENP-N binding

For the nucleosomes assembled on the ASP DNA presented above, we observed a significant impact of CENP-N binding on CENP-A-DNA contacts. Also, both CENP-A and H3 nucleosomes in unbounded state demonstrated highly similar histone core-DNA contacts (see Fig. 5b). Assembly of histones on the Widom 601 DNA revealed a different behavior: the average number of contacts of H3 within SHL 2 to 3.5 decreased from around 63 to 38 (see Fig. S16a), accompanied by substantially larger fluctuations in contact occupancy during the CG-MD simulations (see Fig. S8 and S16). In contrast, the unbound centromeric nucleosome exhibits a smaller difference in the average number of contacts within the same region, with average values of 51 and 59 for Widom 601 and ASP DNA, respectively. These observations indicate that the replacement of ASP with Widom 601 has a more pronounced effect on local H3-DNA interactions than on the corresponding CENP-A-DNA interactions. Thus, the difference between canonical and centromeric nucleosomes is greater with Widom 601 than ASP in terms of average histone-DNA contact numbers. However, analysis of the DNA base-pair-step parameters (see Fig.7a-f) reveals the opposite trend in the extent of local geometrical differences between DNA. Differences occur at positions A-T (89) and A-T (90) and then extend from positions G-C (93) to C-G (108). The strongest coupled changes are concentrated between positions A-T (96) and A-T (102), particularly within the G-C rich segment spanning positions G-C (99) to G-C (101). At position G-C (99), pronounced differences are observed in rise, slide, roll, tilt, and twist, while positions G-C (100) and G-C (101) show substantial changes in shift, slide, tilt, and twist. Toward the end of the analyzed region, from positions T-A (103) to C-G (108), the differences become predominantly associated with roll (see Fig.7f). Compared with ASP DNA, the main region of canonical-centromeric divergence shifts from positions T-A (93)-T-A (97) to positions A-T (96) to A-T (102) with Widom 601 DNA. Overall, Widom 601 DNA exhibits fewer individual inter-base pair parameter deviations than ASP DNA. In particular, the numbers of rise, tilt, and twist deviations decrease, whereas the number of roll deviations remains comparable and extends further toward the end of the analyzed region. Thus, ASP DNA produces more extensive, simultaneous changes in multiple translational and rotational parameters near the beginning of the SHL 2 to 3.5 region. Widom 601 redistributes the principal deformation toward the G-C rich segment and produces predominantly roll-associated differences at the later positions.

The analysis of changes upon the CENP-N binding revealed a different impact on the histone-DNA contacts in centromeric nucleosomes in the presence of both sequences. It reduces the average number of CENP-A-DNA contacts only by 19.6% (from 51 to 41) with Widom 601 DNA, but has a substantially larger reduction by 47.5% (from 59 to 31) in the ASP DNA. The residue-wise DNA contact occupancy, 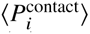 analysis (see Fig. S16 c-d) indicates that Q89 of CENP-A shows the largest decrease in DNA-contact occupancy upon CENP-N binding, while N85, W86 and Q87 retain high DNA-contact occupancies using Widom 601. This suggests the absence of sequence-dependent reorganization of the L1-loop neighboring amino acids observed in ASP. Furthermore, contacts of F84 and Q89 with DNA are more pronounced in Widom 601 than we observed in ASP (see Fig. S8 and S14). In general, analyses show that changes in the binding interface responsible for CENP-N binding are significantly smaller when centromeric nucleosomes assemble on Widom 601. However, similarly as we explained above, changes in the histone-DNA contact occupancy do not directly follow geometrical differences of DNA possessing different sequences. In Fig. 7, we observe that CENP-N binding produces a more spatially extensive change in DNA geometry with Widom 601 than with ASP DNA (Fig. 6). In the ASP system, the principal changes are concentrated at positions T-A (94) to G-C (101), with the strongest coupled translational and rotational changes occurring at positions T-A (94) to T-A (97). In contrast, the Widom 601 system shows deviations across nearly the entire analyzed region, from positions A-T (89) to C-G (108), with prominent changes at positions A-T (96), T-A (97), G-C (100) to T-A (103), and C-G (108). The increase is particularly pronounced for roll and twist, indicating that rotational rearrangements dominate the response to CENP-N binding in the Widom 601 system. With ASP DNA, the CENP-N-induced changes are concentrated mainly near the beginning of the SHL +2 to +3.5 region. In contrast, with Widom 601 DNA, the changes extend to the end of this region. For example, at the C-G step at position 108, CENP-N binding produces a pronounced 34.86^◦^ decrease in twist (Fig. 7f) together with a 3.03 Å increase in shift (Fig. 7a). Thus, Widom 601 reduces the loss of histone-DNA contacts upon CENP-N binding but distributes the accompanying changes in DNA geometry across a broader region. Together, these observations suggest that, although CENP-N recognizes CENP-A nucleosomes largely independently of DNA sequence, as known in the literature (62), the local structural rearrangements associated with its binding are modulated by the underlying DNA sequence.

**Figure 7:**
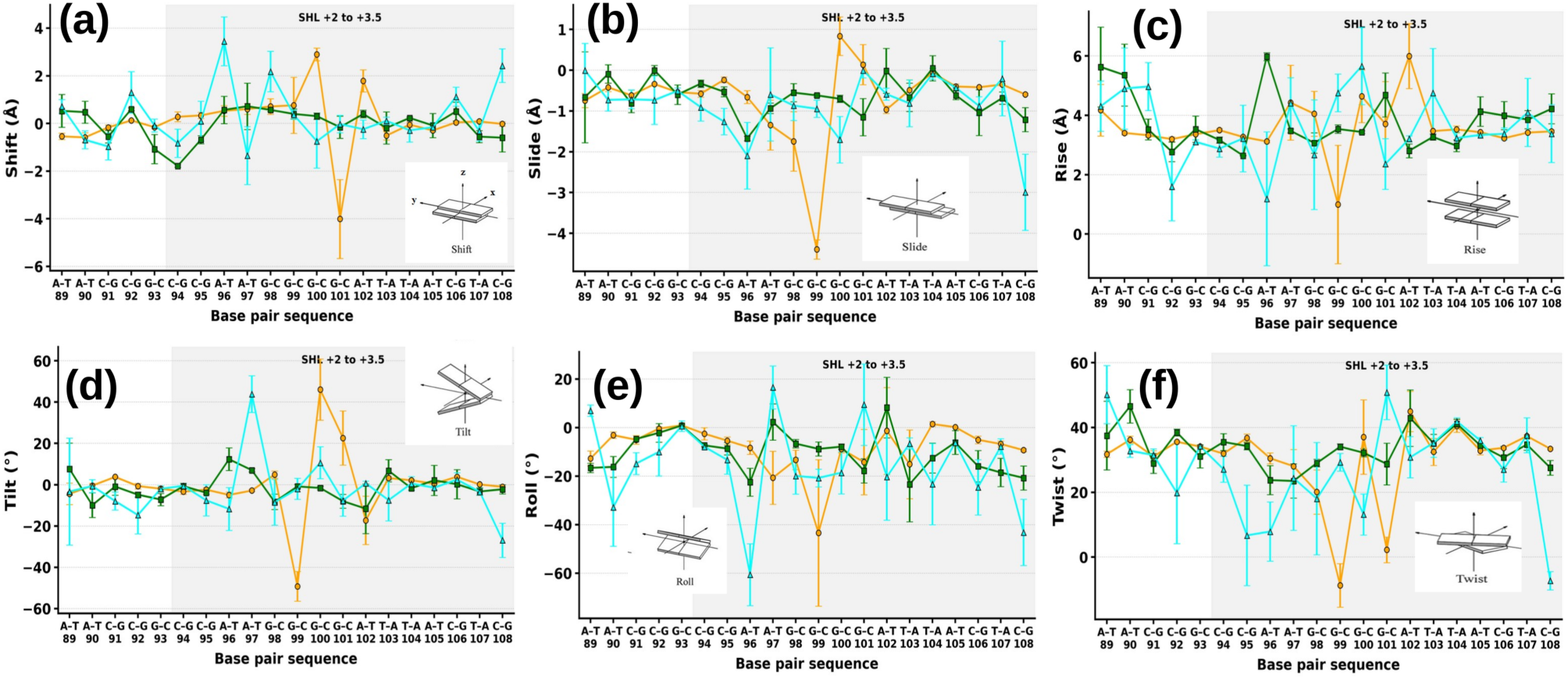
Changes in the DNA inter-base-pair parameters Shift (a), Slide (b), Rise (c), Tilt (d), Roll (e), and Twist (f) for the different systems, with the SHL 2–3.5 region highlighted. A schematic illustration of each parameter is included in the corresponding panel. All analyses were performed using the Widom 601 DNA sequence. The centromeric and canonical nucleosome systems are represented in green and orange, respectively, while the centromeric nucleosome with CENP-N is represented in cyan. Error bars were calculated from three independent replicas.

## CONCLUSION

In summary, we employed long-timescale coarse-grained molecular dynamics simulations to investigate how histone variant identity and DNA sequence collectively influence nucleosome structure, conformational dynamics, and histone-DNA interactions in both canonical and centromeric nucleosomes. Using the same *α*-satellite-derived DNA sequence as reference, our CG simulations reveal distinct structural and dynamic properties of centromeric and canonical nucleosomes. Compared with canonical H3 nucleosomes, CENP-A nucleosomes exhibit reduced histone-core fluctuations and smaller DNA-gaping distances, together with distinct patterns of histone-tail-DNA interactions networks. CENP-N binding further remodels the centromeric nucleosome by substantially reducing local CENP-A-DNA contacts within SHL 2 to 3.5 and altering the base-pair-step parameters of this DNA region. These structural changes are concentrated mainly near the beginning of the SHL 2 to 3.5 region and are dominated by changes in the rotational parameters, particularly roll and tilt. Thus, CENP-N binding induces a localized reorganization of the CENP-A–DNA interface within the ASP-containing centromeric nucleosome.

Extending the analysis to Widom 601 DNA reveals that the underlying sequence alters different components of nucleosome dynamics in distinct ways. Widom 601 generally reduces histone-core fluctuations in several regions, indicating a more constrained histone core, although some chain- and region-specific increases remain. Widom 601 redistributes histone-tail–DNA interactions, reducing H3 and H4 tail contacts in the canonical nucleosome while increasing CENP-A and H2A C-terminal contacts and decreasing H2B tail contacts in the centromeric nucleosome. Widom 601 has little effect on DNA gaping in the canonical nucleosome but increases the gaping distance in the centromeric system, thereby reducing the difference between the two nucleosome types. At the local CENP-A-DNA interface, CENP-N binding causes a smaller loss of contacts with Widom 601 than with ASP DNA, but the accompanying changes in DNA base-pair-step parameters extend across a broader region, reaching the end of SHL 2 to 3.5. Thus, the effect of Widom 601 is not simply greater nucleosome stabilization or compaction. Instead, the sequence redistributes local interactions and modifies how the CENP-A-DNA interface accommodates CENP-N binding, consistent with recognition through localized structural adaptation rather than large-scale rearrangement of the nucleosome architecture.

Taken together, our results demonstrate that the effects of DNA sequence on nucleosome organization depend on histone composition. The distinct sequence-dependent responses of canonical and centromeric nucleosomes are reflected in both histone-DNA contact patterns and local DNA geometry, although changes in these structural properties do not necessarily follow the same trends. Furthermore, the DNA sequence differently modulates the local conformational rearrangements of DNA region associated with CENP-N binding. These findings highlight the importance of considering both DNA sequence and histone composition when investigating nucleosome structure and dynamics. Sequence-dependent changes in histone-DNA interactions observed in canonical nucleosomes cannot be directly extrapolated to centromeric nucleosomes. Finally, although CENP-N recognition is largely conserved across different DNA sequences, the associated local conformational rearrangements can differ. These observations are particularly relevant to the interpretation of experimental structures and simulations employing artificial nucleosome-positioning sequences, such as Widom 601, as models of centromeric nucleosomes. Similar sequence-dependent effects may also be relevant to nucleosomes containing other histone variants, where the interplay between DNA sequence and histone composition could influence local structural dynamics and protein interactions. Furthermore, histone modifications, which were not considered in the present study, may introduce an additional layer of regulation by modulating nucleosome structure and associated protein interactions.

## DATA AVAILABILITY

The results of this study are presented in the manuscript and Supporting Material. Molecular dynamics trajectories and simulation analysis scripts are being prepared for deposition in the Molecular Dynamics Data Bank (MDDB) and KIT GitLab, respectively, and will be publicly available no later than publication of the peer-reviewed article. Until then, these materials are available from the corresponding author upon reasonable request.

## AUTHOR CONTRIBUTIONS

Abhik Ghosh Moulick: Writing- review and editing, Writing- original draft, Visualization, Validation, Software, Methodology, Investigation, Formal analysis, Data curation, Conceptualization. Sylvia Erhardt: Writing- review and editing, Validation, Funding acquisition. Mariana Kozlowska: Writing- review and editing, Supervision, Resources, Project administration, Methodology, Funding acquisition, Validation, Formal analysis, Conceptualization.

## ACKNOWLEDGMENTS

This research was made possible by funding from the Carl-Zeiss-Stiftung and Center SynGen. The authors gratefully acknowledge the computing time provided on the high-performance computer HoreKa by the National High-Performance Computing Center at KIT (NHR@KIT). This center is jointly supported by the Federal Ministry of Education and Research and the Ministry of Science, Research and the Arts of Baden-Württemberg, as part of the National High-Performance Computing (NHR) joint funding program (https://www.nhr-verein.de/en/our-partners). HoreKa is partly funded by the German Research Foundation (DFG). We acknowledge support by the KIT-Publication Fund of the Karlsruhe Institute of Technology.

## SUPPORTING MATERIAL

Supporting Material is available online with this article.

