## Supporting Material for "DNA Sequence Differentially Modulates Histone Core and Tail Interactions in Canonical and Centromeric Nucleosomes"

### System preparation and coarse-grained MD simulation

All nucleosome systems were prepared and simulated following earlier studies [1, 2] in GROMACS 2019.3[3]. In brief, all-atom refined structures after modeling using AlphaFold3 were protonated using the PDB2PQR server [4] under neutral pH conditions and with AMBER naming conventions required for SIRAH. The resulting structures were then mapped to a coarse-grained representation using the SIRAH 2.2 tools. The mapped systems were solvated in a periodic boundary condition (PBC) box using SIRAH WT4 water beads [5]. Systems were neutralized and brought to physiological conditions by adding $Na^{+}$ and $Cl^{-}$ ions to achieve a salt concentration of 0.15 M. The number of ions, box dimensions, and number of solvent beads are listed in Table S1. A truncated octahedral simulation box was used for centromeric system while cubic box was used for canonical system, maintaining a minimum distance of 20 Å between the solute and the box boundaries to avoid interactions with periodic images. Energy minimization was performed in two stages: (i) 50,000 steps with side chains minimized while restraining the backbone, and (ii) 5,000 steps of unrestrained minimization of the entire system, using the steepest descent algorithm in both cases. Side-chain equilibration improved structural stability by avoiding distortions of secondary structure elements. Subsequently, solvent equilibration was performed through a 5 ns simulation with harmonic restraints applied to all coarse-grained beads. Temperature was maintained at 300 K using the V-rescale thermostat [6], and pressure was kept at 1 bar using the Parrinello–Rahman barostat [7]. Before production, an additional 10 ns restrained simulation was performed to allow DNA relaxation at the protein–DNA interface. Final production simulations were carried out for 10 $\mu s$ in the absence of restraints, with a 20 fs time step. Long-range electrostatics were treated using the PME method with a 12 Å cutoff and a grid spacing of 2 Å, while van der Waals interactions were also cut off at 12 Å. Three independent replicas were simulated for each system using different random seeds, and the final 5 $\mu$s of each trajectory were used for analysis.

SIRAH-Backmap tools [8] were used to backmap the coarse-grained molecular dynamics trajectories to atomistic resolution. The backmapping procedure reconstructs atomistic positions on a per-residue basis [9], preserving internal geometric coordinates. This is followed by protonation and energy minimization using the ff14SB atomistic force field [10] implemented in the tleap module of AmberTools [11].

**Validation of the CENP-A nucleosome model on Widom 601 DNA**

To validate the structural integrity of the model, we compared our model system of centromeric nucleosome reconstituted on Widom 601 sequence (referred as model system) with deposited cryo-EM structure from PDB ID: 6TEM (referred as deposited). Long timescale CG simulations of 15 $\mu$s were performed for both systems, and several structural parameters, including histone-DNA contacts, radius of gyration ($R_g$), RMSD, and residue-wise RMSF, were analyzed to assess the similarity between the two nucleosome architectures. Fig. S9a-d shows time dependent change of the histone core and DNA contact ($N_C$). Here, we used the CG trajectories to perform contact analysis by considering interactions between the protein backbone GC beads and the DNA backbone PX beads. A cutoff distance of 7 Å was used to define a contact, following previous studies [1]. For the CENP-A histone, the time dependent contact profile exhibits only minor differences between the model and deposited systems, while for the remaining histones the $N_C$ profiles largely overlap, suggesting that that the modeling procedure considered in current study does not significantly perturb the native protein–DNA packing interactions. The other copy of each histone exhibits a similar behavior, with only minor differences between the model and deposited systems (see Fig. S9e-h).

To further assess the structural stability of the model, we computed global structural parameters like RMSD and $Rg$ of DNA. The average values of $Rg$ and RMSD, represented in Fig. S10a, remain largely unchanged between the two systems, indicating that the overall nucleosome architecture is preserved upon modeling. We further quantified local structural deviations by calculating the per base pair difference in RMSF values ($\Delta RMSF$). Fig. S10b indicates minimal differences within the nucleosomal core region. Overall, these results demonstrate that the structural integrity and native protein–DNA organization of the deposited CENP-A nucleosome are well preserved in the reconstructed model, supporting its suitability for subsequent analyses of sequence dependent nucleosome dynamics.

Table S1: Summary of the systems used for CG-MD simulations with SIRAH, including the simulation box dimensions, number of coarse-grained beads, solvent beads, and $Na^+$ and $Cl^-$ ions.

| **System** | **DNA** | **PBC Box (Å)** | **Beads** | **Solvent** | **$Na^+$** | **$Cl^-$** |
|---|---|---|---|---|---|---|
| Centromeric nucleosome | ASP | 171.3 × 161.5 × 139.8 | 48078 | 10222 | 399 | 247 |
| Centromeric nucleosome | Widom 601 | 171.3 × 161.5 × 140.0 | 48286 | 10279 | 399 | 251 |
| Centromeric nucleosome + CENP-N | ASP | 192.1 × 181.3 × 157.0 | 67454 | 14620 | 536 | 387 |
| Centromeric nucleosome + CENP-N | Widom 601 | 192.3 × 181.3 × 157.0 | 67676 | 14681 | 535 | 390 |
| Canonical nucleosome | ASP | 210 × 210 × 210 | 114459 | 26587 | 896 | 930 |
| Canonical nucleosome | Widom 601 | 213 × 213 × 213 | 117837 | 27448 | 783 | 805 |
| Centromeric nucleosome (deposited structure) | Widom 601 | 150.8 × 142.2 × 123.1 | 32319 | 6708 | 309 | 114 |

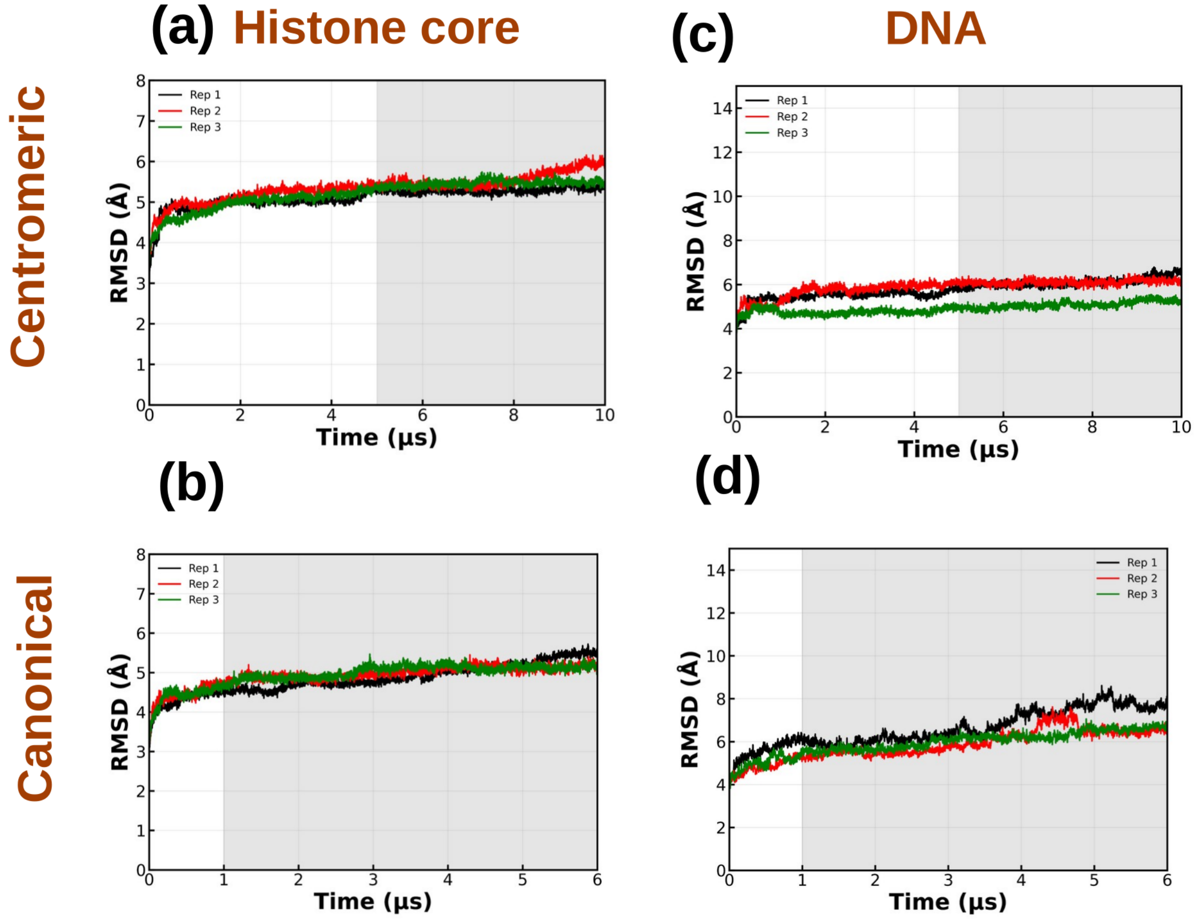


Fig. S1: RMSD of the histone core for the (a) centromeric and (b) canonical nucleosome systems across different replicas based on the ASP sequence. The canonical nucleosome simulations were taken from our previously published work[12]. Histone tails were excluded from the analysis because of their inherent flexibility. Panels (c) and (d) show the RMSD of DNA for the centromeric and canonical systems, respectively. The last 5 $\mu$s of each trajectory, used for the subsequent analyses, are highlighted in grey.

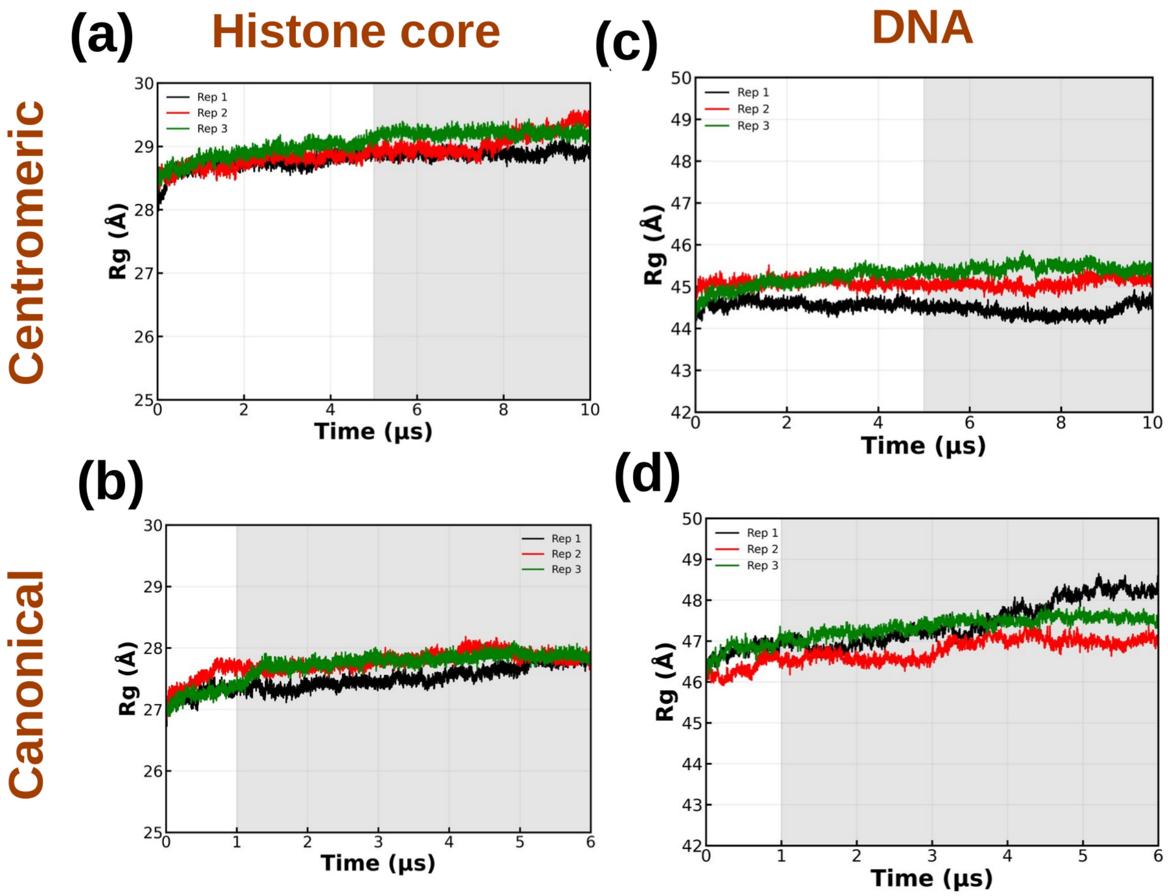


Fig. S2: Radius of gyration ($R_g$) of the histone core for the (a) canonical and (b) centromeric nucleosome systems across different replicas based on the ASP sequence. Panels (c) and (d) show the RMSD of DNA in the canonical and centromeric systems, respectively. The last 5 $\mu$s of each trajectory, which were used for subsequent analyses, are highlighted in grey.

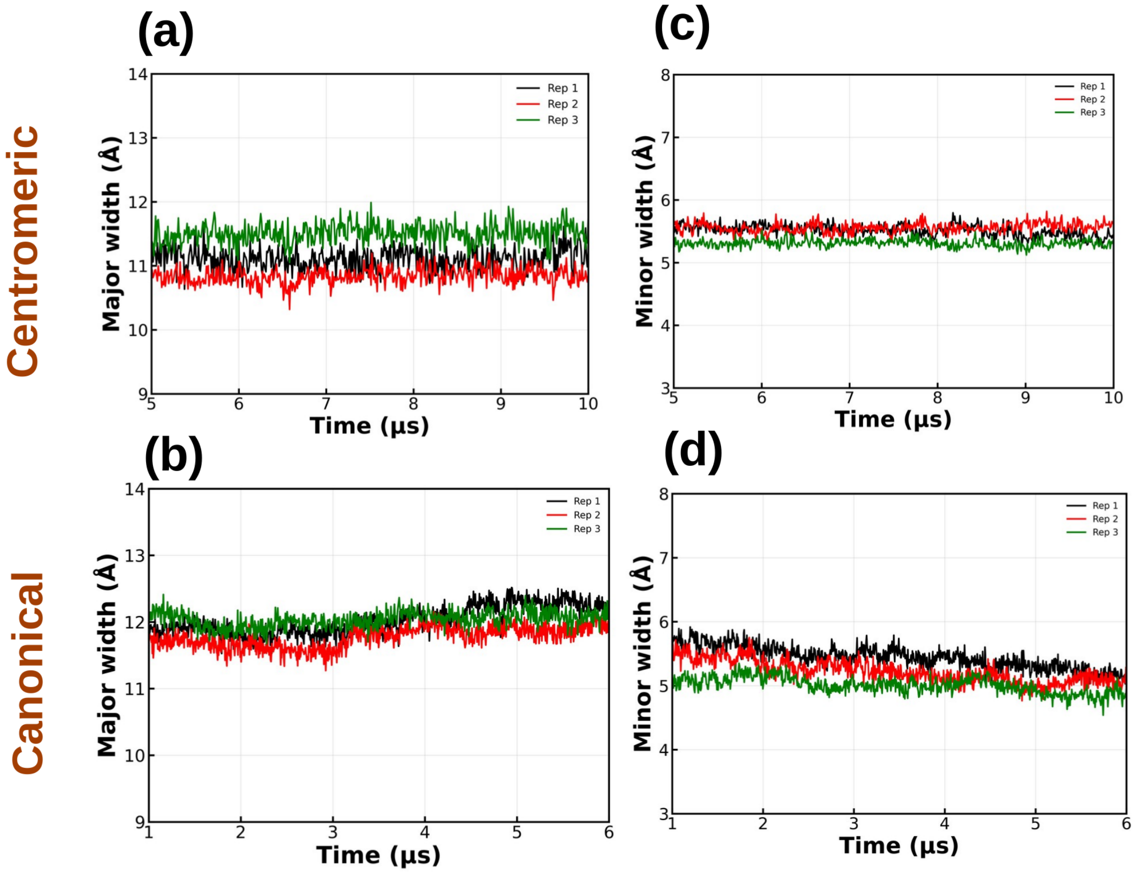


Fig. S3: DNA major groove width for the (a) centromeric and (b) canonical nucleosome systems across different replicas. Panels (c) and (d) show the time dependent change of minor groove width of the DNA in the centromeric and canonical systems, respectively. For each system, the last 5 $\mu$s of the trajectory are depicted.

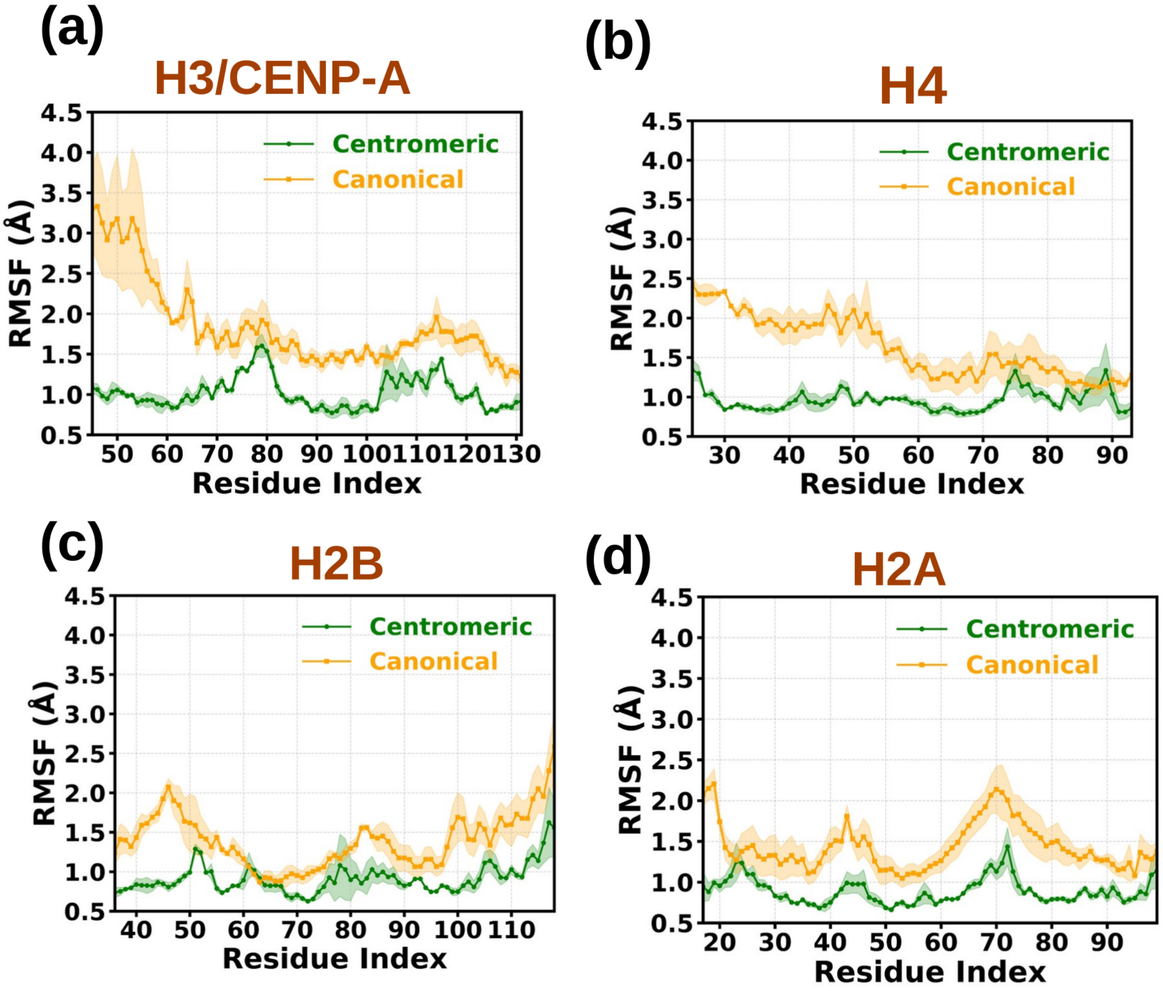


Fig. S4: Root mean square fluctuations (RMSF) of the histone core domains of the second copy of the histones: (a) H3 (canonical) or CENP-A (centromeric), (b) H4, (c) H2A, and (d) H2B. Orange and green correspond to the canonical and centromeric nucleosome systems, respectively. The RMSF profiles were obtained by averaging over three independent simulation replicas for each system. Solid lines represent the mean RMSF values, while the shaded regions indicate the corresponding standard errors.

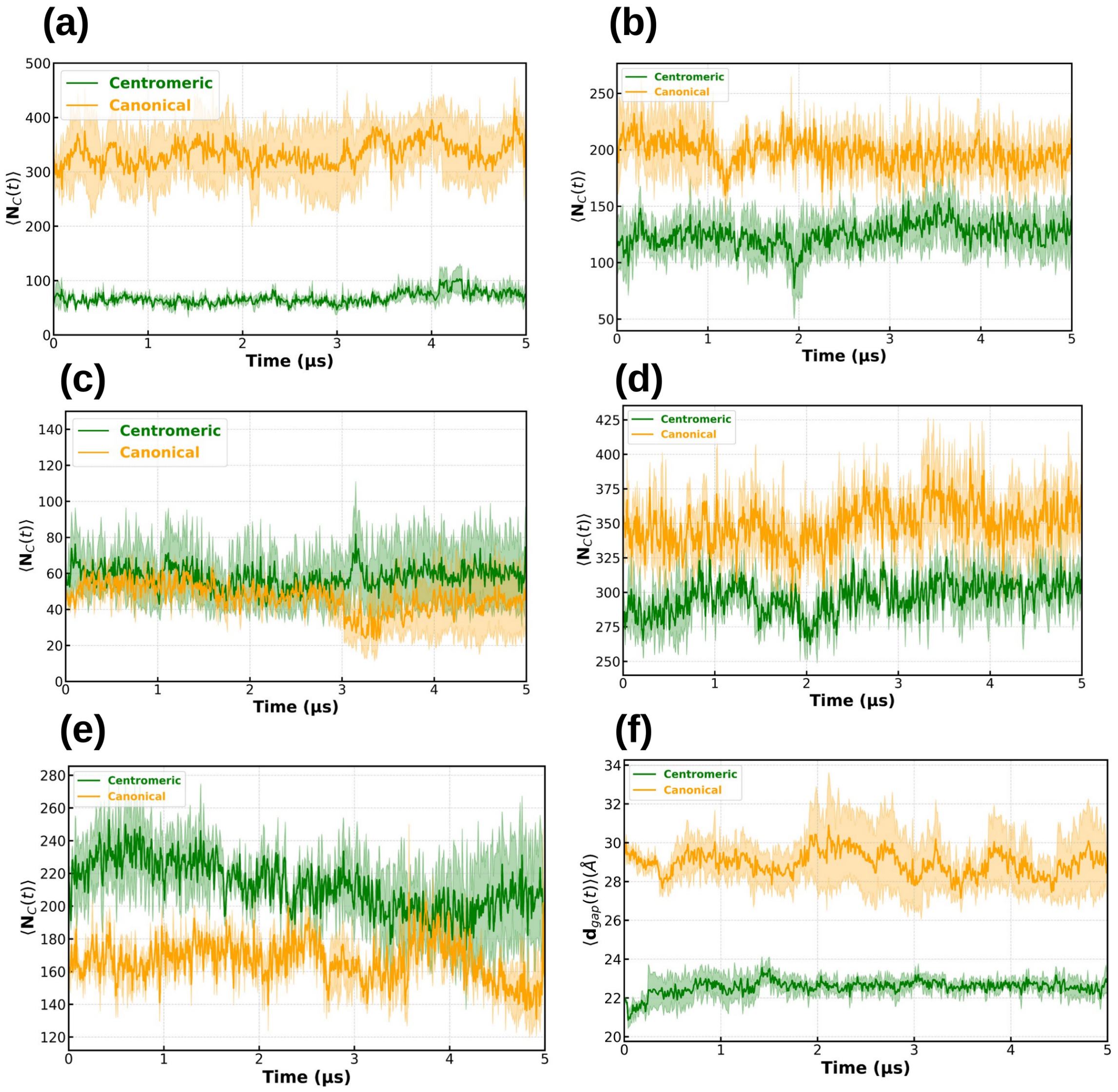


Fig. S5: Time evolution of the the average number of contacts ($\langle N_C(t) \rangle$) during the simulation between different histone tails and ASP DNA for the canonical (orange) and centromeric (green) nucleosome systems. Panels show the (a) H3/CENP-A N-terminal, (b) H4 N-terminal, (c) H2A C-terminal, (d) H2B N-terminal, and (e) H2A N-terminal tails. Panel (f) shows change of the DNA gaping distance ($\langle d(t) \rangle$). The average values were obtained by averaging over the last 5 $\mu$s of each of the three independent simulation replicas for each system. Solid lines represent the average values, while the shaded regions indicate the corresponding standard errors.

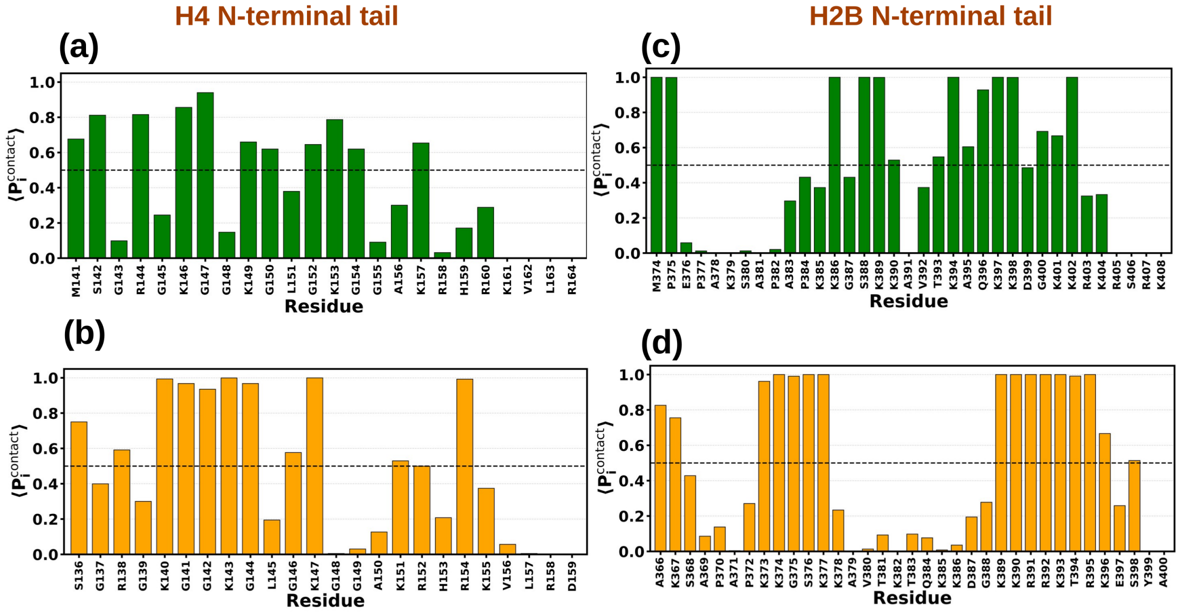


Fig. S6: Residue-wise average DNA-contact occupancy, $\langle P_i^{\text{contact}} \rangle$, for the H4 N-terminal tail in the (a) centromeric and (b) canonical systems. Panels (c) and (d) show the corresponding DNA-contact occupancies for the H2B N-terminal tail in the centromeric and canonical systems, respectively. Residues with $\langle P_i^{\text{contact}} \rangle \geq 0.5$ were considered persistently DNA-contacting residues.

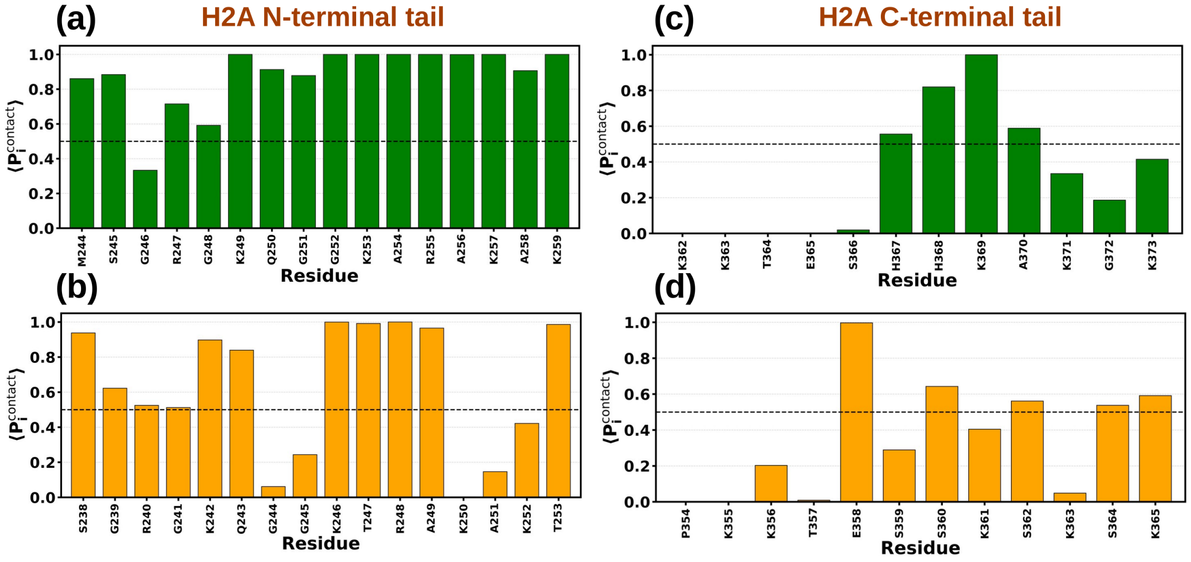


Fig. S7: Residue-wise average DNA-contact occupancy, $\langle P_i^{\text{contact}} \rangle$, for the H2A N-terminal tail in the (a) centromeric and (b) canonical systems. Panels (c) and (d) show the corresponding DNA-contact occupancies for the H2A C-terminal tail in the centromeric and canonical systems, respectively.

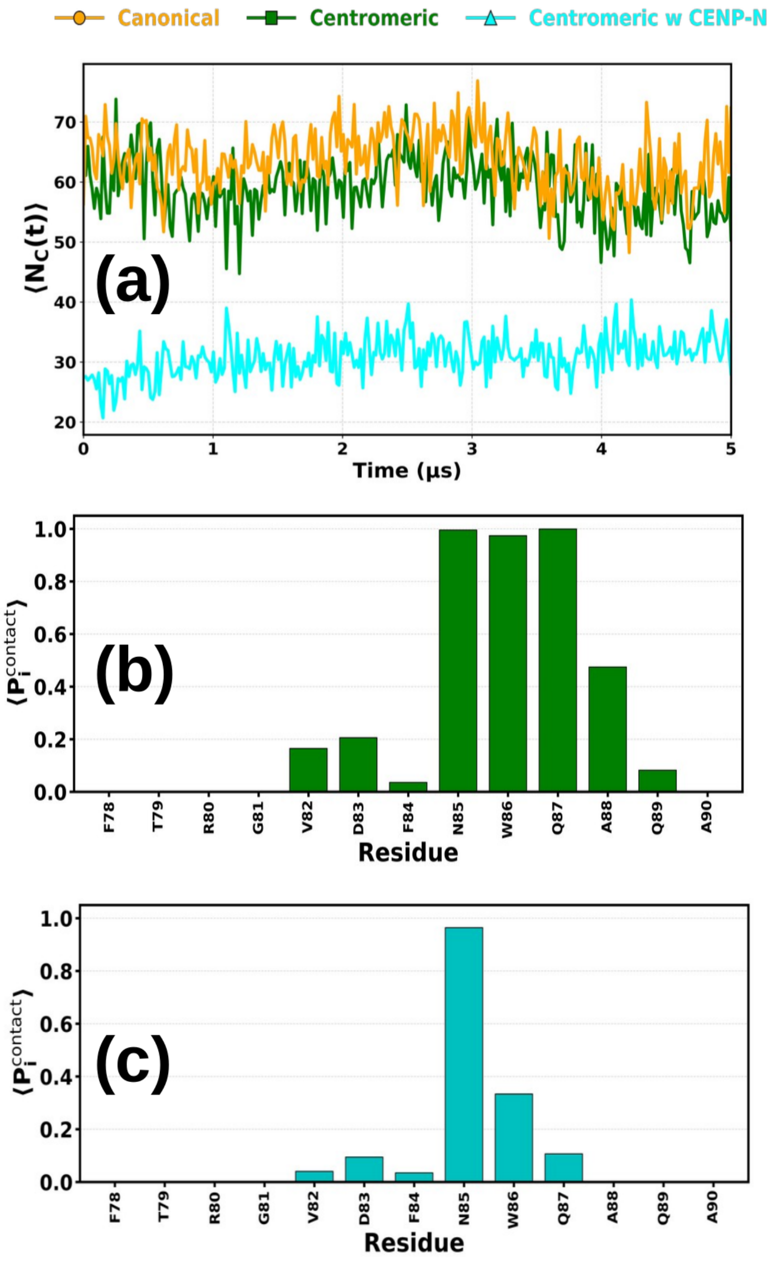


Fig. S8: (a) Time evolution of CENP-A/H3 DNA contacts within the SHL+2 to SHL+3.5 region corresponding to the CENP-N binding interface. The comparison includes (i) canonical nucleosome (orange), (ii) centromeric nucleosome (green), and (iii) centromeric nucleosome upon CENP-N binding (sky) on the ASP sequence. Residue-wise average DNA-contact occupancy, $\langle P_i^{\mathrm{contact}} \rangle$, of the residues in the L1 loop and neighboring residues of the CENP-A histone in the absence (b) and presence (c) of CENP-N.

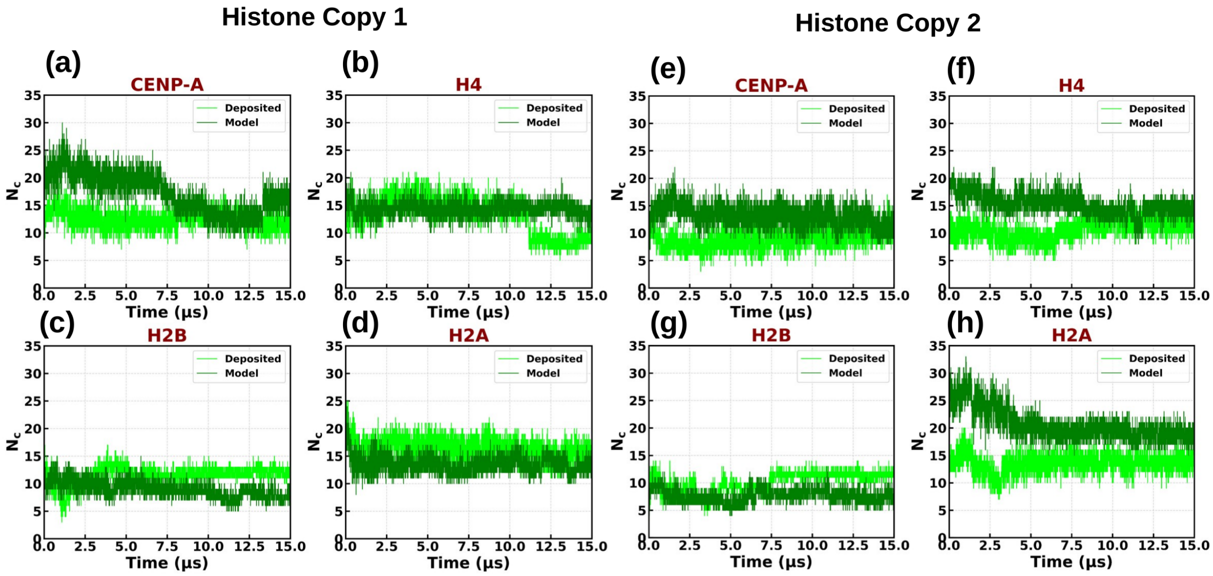


Fig. S9: Time evolution of histone core–DNA contacts ($N_C$) for CENP-A, H4, H2B, and H2A. The deposited structure corresponds to PDB ID 6TEM, whereas the model structure was generated by replacing the ASP DNA sequence in the centromeric nucleosome model with the Widom 601 DNA sequence while retaining the histone core. Panels (a)–(d) show the changes in $N_C$ for histone copy 1, whereas panels (e)–(h) show the corresponding changes for histone copy 2.

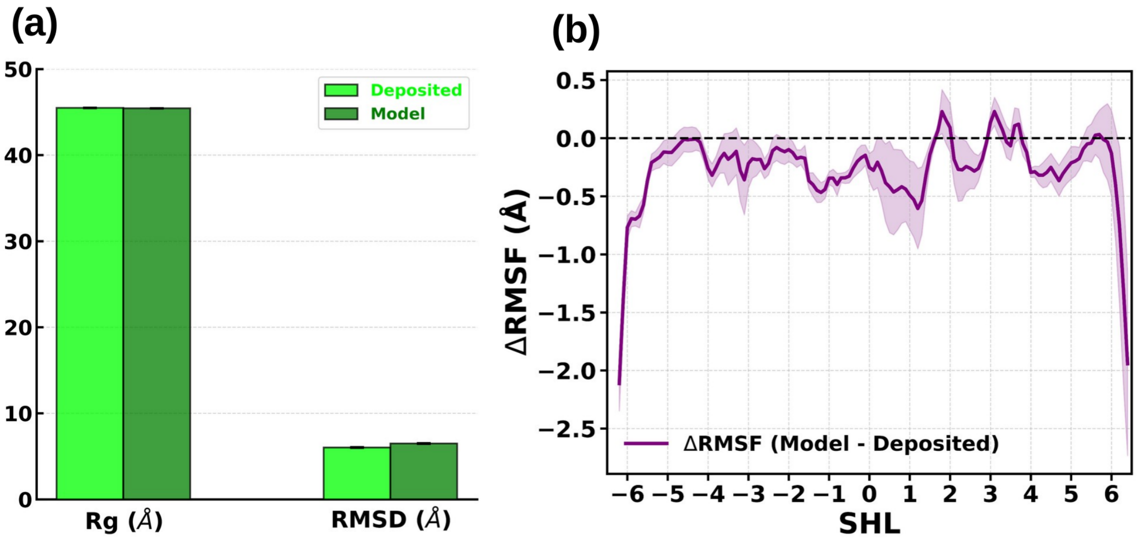


Fig. S10: (a) Bar plot showing the average radius of gyration (Rg) and root-mean-square deviation (RMSD) for the deposited and modeled structures. The modeled structure is shown in deep green, while the deposited structure is shown in light green. (b) Per-base-pair RMSF differences between the modeled and deposited DNA represented in superhelical location (SHL) notation.

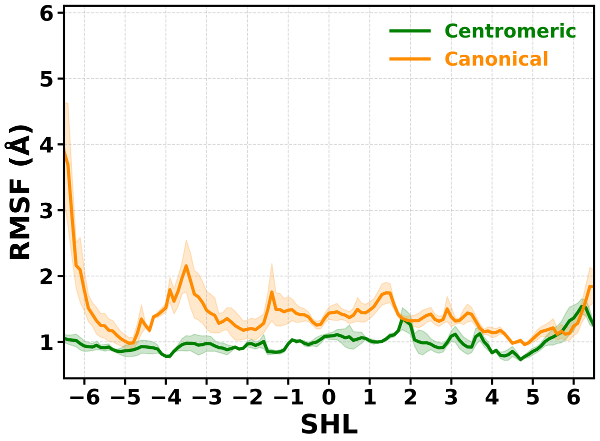


Fig. S11: Comparison of DNA root-mean-square fluctuations (RMSF) between centromeric (green) and canonical (orange) nucleosomes assembled on Widom-601 DNA. RMSF values are plotted as a function of superhelical location (SHL). Solid lines represent the mean RMSF calculated from three independent simulation replicas, while shaded regions indicate the standard error of the mean.

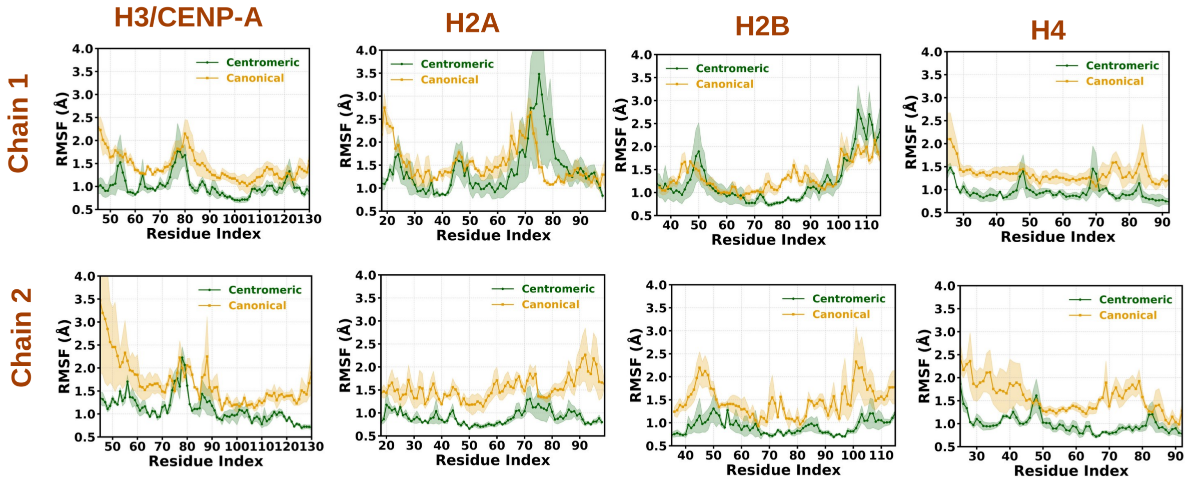


Fig. S12: Root mean square fluctuations (RMSF) of the histone core domains of the both chains (Chain 1 and Chain 2) of the histones on Widom 601 sequence: H3 (canonical) or CENP-A (centromeric), H4, H2A, and H2B. Orange and green correspond to the canonical and centromeric nucleosome systems, respectively. The RMSF profiles were obtained by averaging over three independent simulation replicas for each system. Solid lines represent the mean RMSF values, while the shaded regions indicate the corresponding standard errors.

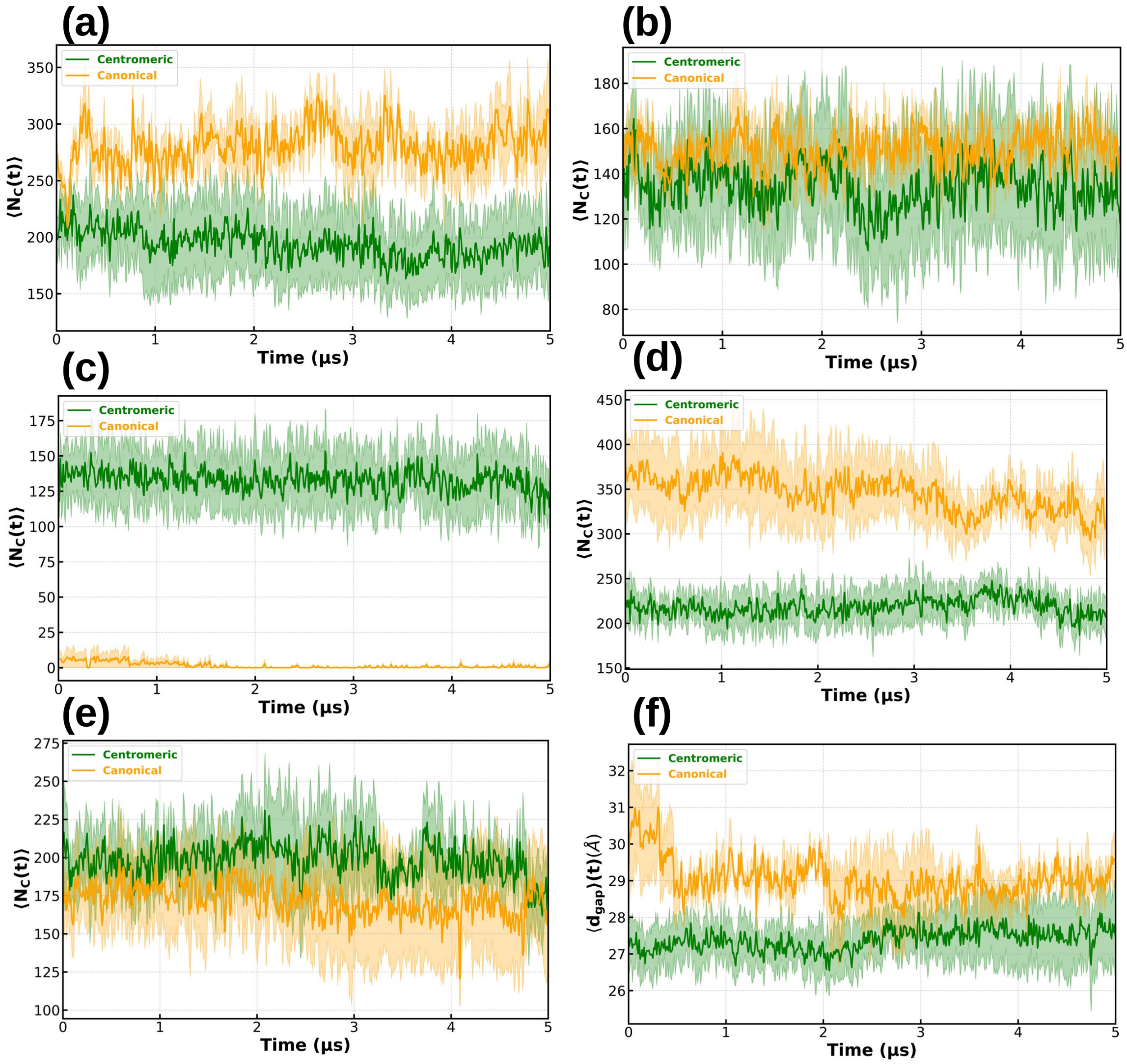


Fig. S13: Time evolution of the average number of contacts ($\langle N_C(t)\rangle$) between different histone tails and Widom 601 DNA for the centromeric (green) and canonical (orange) nucleosome systems. Panels show the (a) CENP-A/H3 N-terminal, (b) H4 N-terminal, (c) H2A C-terminal, (d) H2B N-terminal, and (e) H2A N-terminal tails. Panel (f) shows the time evolution change in the DNA gaping distance ($\langle d(t)\rangle$). The average values were obtained by averaging over three independent simulation replicas for each system. Solid lines represent the average values, while the shaded regions indicate the corresponding standard error of the mean across replicas.

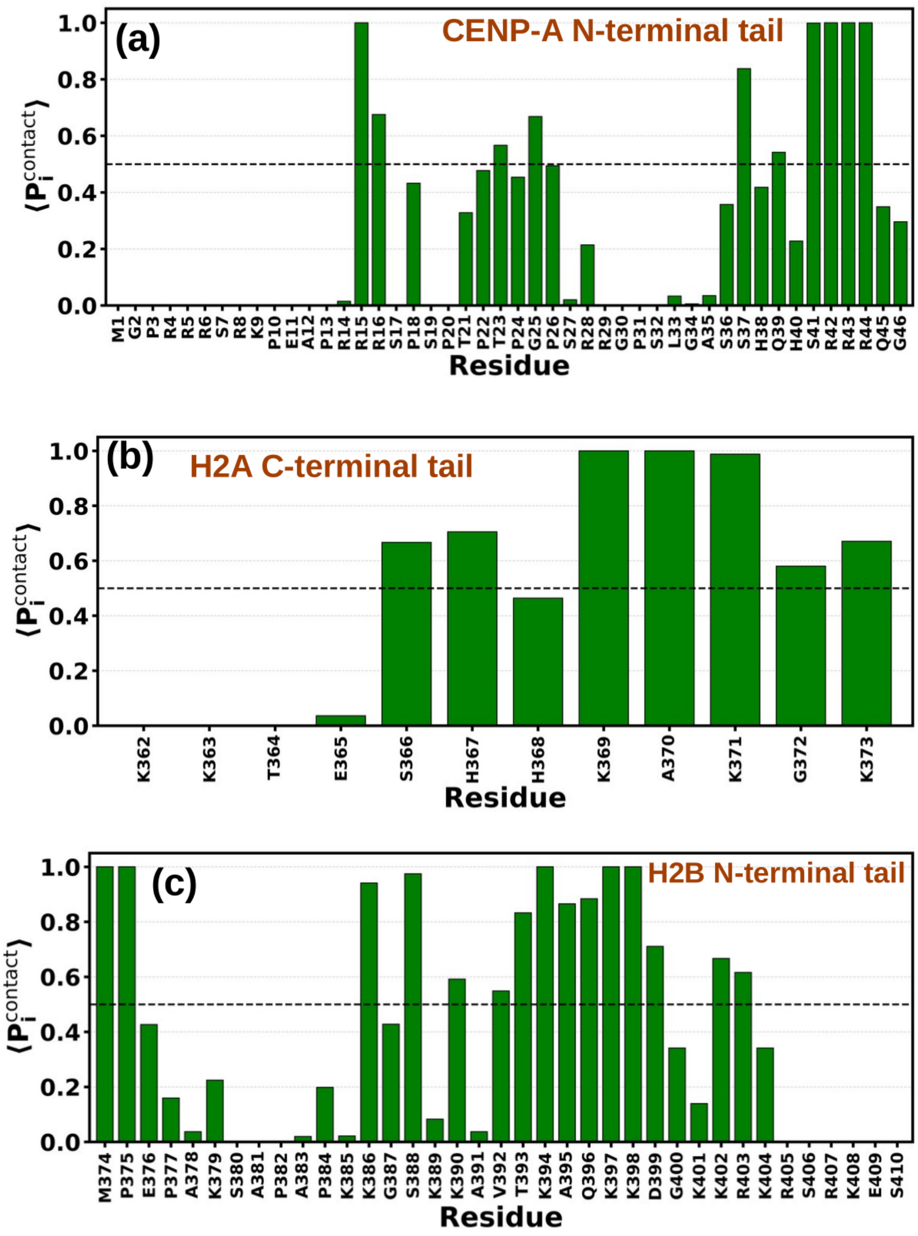


Fig. S14: (a) Residue-wise average DNA-contact occupancy, $\langle P_i^{\text{contact}} \rangle$, for the CENP-A N-terminal tail in the centromeric system for Widom 601 sequence. Panels (b) and (c) show the corresponding DNA-contact occupancies for the H2A C-terminal tail and H2B N-terminal tail respectively. Residues with $\langle P_i^{\text{contact}} \rangle \geq 0.5$ were considered persistently DNA-contacting residues.

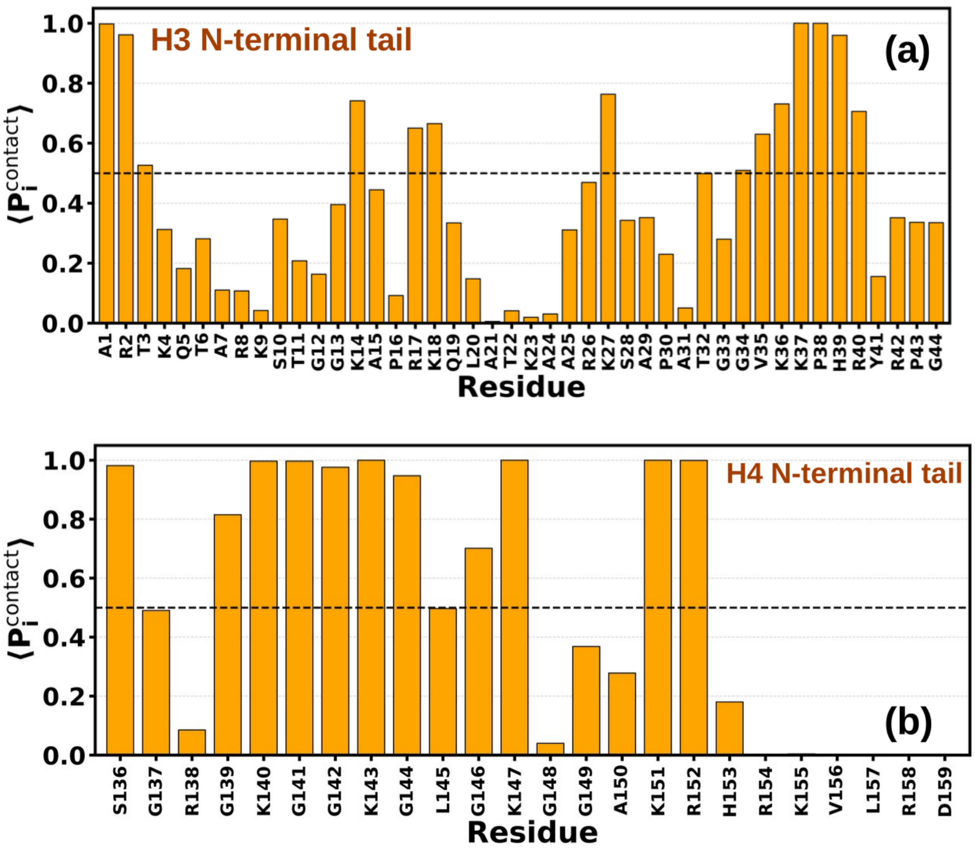


Fig. S15: ((a) Residue-wise average DNA-contact occupancy, $\langle P_i^{\mathrm{contact}} \rangle$, for the H3 N-terminal tail in the canonical system for Widom 601 sequence. Panels (b) show the corresponding DNA-contact occupancies for the H4 N-terminal. Residues with $\langle P_i^{\mathrm{contact}} \rangle \geq 0.5$ were considered persistently DNA-contacting residues.

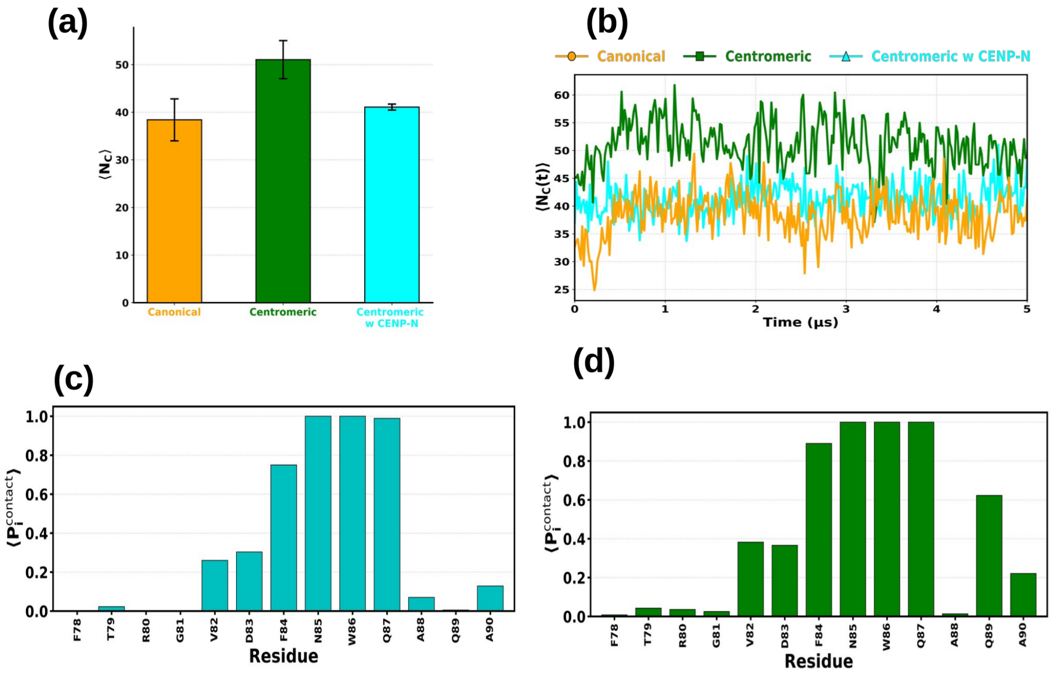


Fig. S16: (a) Bar plots showing the average number of contacts ($\langle N_c \rangle$) between the SHL +2 to +3.5 region and CENP-A in the centromeric nucleosome (green) or H3 in the canonical nucleosome (orange). The average number of contacts decreases upon CENP-N binding to the centromeric nucleosome (cyan). (b) Time evolution of CENP-A/H3 DNA contacts within the SHL+2 to SHL+3.5 region corresponding to the CENP-N binding interface. The comparison includes (i) canonical nucleosome (orange), (ii) centromeric nucleosome (green), and (iii) centromeric nucleosome upon CENP-N binding (sky) on Widom 601 sequence. Residue-wise average DNA-contact occupancy, $\langle P_i^{\text{contact}} \rangle$, of the residues in the L1 loop and neighboring residues of the CENP-A histone in the absence (c) and presence (d) of CENP-N.
